# Graphene nanoflakes for acute manipulation of membrane cholesterol and transmembrane signaling

**DOI:** 10.1101/581942

**Authors:** Kristina E. Kitko, Tu Hong, Roman M. Lazarenko, J. Rafael Montenegro-Burke, Amy T. Shah, Yuchen Zhang, Krishnan Raghunathan, Anne K. Kenworthy, Melissa C. Skala, A. McLean, Ya-Qiong Xu, Qi Zhang

## Abstract

Cholesterol is one of the most essential lipids in eukaryotic cell membranes. However, acute and selective manipulation of membrane cholesterol remains challenging. Here, we report that graphene nanoflakes (GNFs) insert into the plasma membrane and directly interact with cholesterol, resulting in acute cholesterol enrichment – and thus structural and functional changes. Using two representative cell preparations, we explore the utility of GNFs in modifying cell communication pathways sensitive to membrane cholesterol. In fibroblasts, GNFs enhance ATP-induced intracellular Ca^2+^-release by allosteric facilitation of P2Y receptors, a subtype of G protein-coupled receptors, in a cholesterol-dependent manner. In neurons, which possess higher membrane cholesterol levels than most cell types, GNFs further increase cholesterol. Consequently, GNFs change membrane fluidity, especially at synaptic boutons, and potentiate neurotransmitter release by accelerating synaptic vesicle turnover. Together, our results provide a molecular explanation for graphene’s cellular impacts and demonstrate its potential for membrane-oriented engineering of cell signaling.

## Introduction

In most eukaryotic cells, the plasma membrane is the largest structure and the only interface between the intracellular and extracellular environments. Most intercellular communications are transduced through the plasma membrane via integral proteins or the release and uptake of signaling molecules. Therefore, transmembrane proteins, like receptors and ion channels, and exo-/endocytosis machinery, like the SNARE complex, are the focus of basic as well as preclinical investigation and thus the targets for drug discovery. In contrast, lipids have long been treated as structural support for proteins (Singer and Nicolson, 1972). However, new evidence has suggested that lipids can actively regulate signaling proteins by defining their surface localization (Lingwood and Simons, 2010; Simons and Toomre, 2000) or even via direct modification (Hannun and Obeid, 2008; Yeagle, 2014). One of the best examples is cholesterol, a major sterol molecule with a tetracyclic ring structure (Simons and Ehehalt; Yeagle, 1985). Its hydroxyl head can form hydrogen bonds with the carbonyl oxygen of phospholipid head groups, while its hydrocarbon tail wedges into the non-polar core of the bilayer, “gluing” various acyl chains. As such, cholesterol is essential for the formation and stabilization of membrane nanodomains, in which lipids are highly packed and signaling proteins are compartmentalized (Lingwood and Simons, 2010; Pike, 2006; Simons and Toomre, 2000). Recent structural studies have shown that cholesterol can stabilize G protein-coupled receptors or promote their association via direct binding (Cherezov et al., 2007; Hanson et al., 2008). Cholesterol is also involved in Clathrin-dependent and -independent endocytosis (Subtil et al., 1999) and forms the membrane force foci for mechanical sensing (Anishkin and Kung, 2013). Therefore, cholesterol influences major transmembrane signaling pathways.

While artificial liposomes, like giant unilamellar vesicles, are suitable for biophysical investigation, live cell preparations are preferred for studying the physiological and pathological roles of membrane cholesterol (Simons and Gerl, 2010). However, acute and selective manipulation of cholesterol within the plasma membrane of live cells has proven technically challenging (Simons and Gerl, 2010). To date, the most widely used tool is methyl-β-cyclodextrin (MβCD), whose hydrophobic cavity can bind and extract cholesterol from the plasma membrane (Zidovetzki and Levitan, 2007). Statins, a class of HMG-CoA reductase inhibitors, block cholesterol synthesis and chronically reduce cholesterol across the cell including the plasma membranes (Liao and Laufs, 2005). Genetic mutations that disrupt cholesterol transportation, for example NPC1 of Niemann Pick disease, also cause chronic deprivation of cell membrane cholesterol (Vance and Karten, 2014). However, none of these methods are capable of manipulating cholesterol in membrane nanodomains or even in subregions of the plasma membrane (Liao and Laufs, 2005; Vance and Karten, 2014; Zidovetzki and Levitan, 2007). While the ability to increase membrane cholesterol levels would be equally essential to the investigation of its functional contribution, the means to do so are even more limited (Lingwood and Simons, 2010; Zidovetzki and Levitan, 2007). Cholesterol-saturated MβCD is often applied as it releases cholesterol in the presence of a concentration gradient. However, neither the amount nor the rate of such cholesterol release is predictable (Zidovetzki and Levitan, 2007), and this method is less effective for cells with high membrane cholesterol content (Zidovetzki and Levitan, 2007).

To address the challenge of manipulating membrane cholesterol, we turned to carbon nanomaterials, whose surfaces are often hydrophobic(Li et al., 2013b) and thus favorable for lipid interaction. Graphene, a single-layer carbon crystal (Novoselov et al., 2004), emerged as a promising candidate. Computational modeling and electron microscopy studies have suggested that graphene nanoflakes (GNFs), with their sharp edges and hydrophobic surfaces, readily insert into the lipid bilayer, extracting phospholipids and causing the breakdown of prokaryotic cell membranes (Li et al., 2013a; Tu et al., 2013). Eukaryotic cells, however, seem to be spared from this destruction (Bendali et al., 2013; Bramini et al., 2016; Chen et al., 2012; Li et al., 2011; Sahni et al., 2013; Wang et al., 2012). Recent computational studies have suggested that cholesterol, unique to eukaryotic membranes, tightly surrounds inserted GNFs(Zhang et al., 2016), suggesting a possible explanation for the observed differences. These studies raise the possibility of using GNFs as a nanoscale tool to alter membrane cholesterol localization, and, likely, to modify transmembrane signaling. We empirically tested whether GNFs interact with cholesterol in mammalian cell membranes, how this interaction affects cholesterol distribution, and importantly, what then were the functional consequences.

## Results

### Production and characterization of graphene nanoflakes

To obtain high quality GNFs, we used liquid-phase exfoliation (LPE) of graphite powder, which produces single- and few-layer GNFs in a scalable fashion (Geim, 2009; Hernandez et al., 2008). We suspended GNFs uniformly in H_2_O containing 2 wt% polyvinylpyrrolidone (PVP), which prevented aggregation (Fig. 1a). Based on its absorption coefficient at 660nm (Hernandez et al., 2008), we estimated that the concentration of GNFs suspension was about 26 mg/L. To characterize the GNFs, we used Raman spectroscopy (Fig. 1b). At 532 nm excitation, GNFs suspensions exhibited characteristic G peaks (~1,580 cm^−1^) and 2D peaks (~2,700 cm^−1^) similar to bulk graphite (Hernandez et al., 2008). However, unlike graphite, Raman D peaks (~1,350 cm^−1^) were larger in the GNFs suspensions, suggesting the strong edge effects of nanometer-size GNFs (Ferrari et al., 2006). Next, we performed transmission electron microscopy (TEM) to estimate the size and thickness of the GNFs. As shown in Figure 1c, the smooth planar structures and uniform flake edges support the notion that these GNFs are one to few layers with a lateral dimension of a few hundred nanometers. Notably, we did not observe any contamination to the GNFs suspension resulting from the LPE process. Furthermore, we conducted atomic force microscopy (AFM) to measure the size and thickness of the GNFs (Fig. 1d). The majority of the flakes are 1-2 nm thick (Figure 1e&f), which is expected given the tendency of GNFs to aggregate during AFM preparation (Lotya et al., 2009) and the presence of residual surfactants between GNFs and the substrate (Hernandez et al., 2008). Importantly, the distributions of GNFs thickness and size (Fig. 1f) were consistent with previous reports (Hernandez et al., 2008; Lotya et al., 2009).

**Figure 1.**
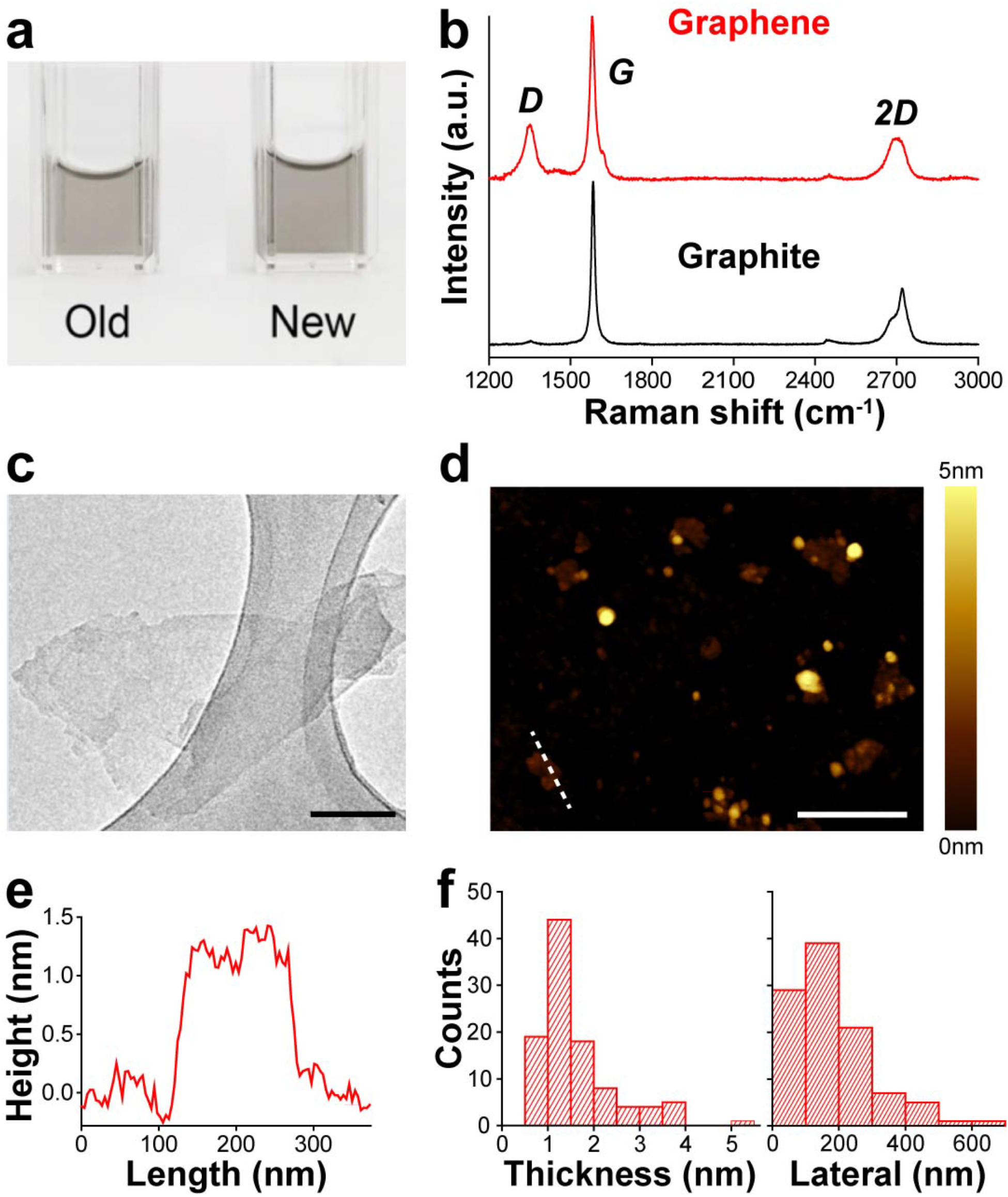
Characterization of graphene nanoflakes. **(a)** Pictures of GNF stock solutions. The left is one-year old and the right is freshly prepared. Both contain 26 mg/L GNFs. **(b)** Raman spectra for GNFs suspended in 2 wt% PVP (red) and bulk graphite (black), respectively. **(c)** Sample TEM image of a GNF. Scale bar, 100 nm. **(d)** Sample AFM image of dispersed GNFs. Scale bar, 500 nm. **(e)** Line profile of a GNF along the white dashed line in **(d). (f)** The distributions of GNF thickness and lateral dimension obtained from AFM image analysis. GNF, graphene nanoflake; PVP, polyvinylpyrrolidone; TEM, transmission electron microscopy; AFM, atomic force microscopy.

### Graphene Directly and Selectively Interacts With Cholesterol

To experimentally validate the graphene–cholesterol interaction, we began by mixing GNF suspension with cell culture media (Fig. 2a). After 1-hour incubation, we isolated GNFs using size-exclusion filtration and measured the amount of cholesterol in each fraction with a quantitative enzymatic assay (see online methods). We found that cholesterol was enriched in the GNFs fraction and depleted in the GNFs-free fraction (Fig. 2a), indicating that GNFs adsorb cholesterol. To determine if cholesterol directly interacted with the GNFs, we took advantage of a well-documented property of graphene – it acts as a broad-spectrum acceptor in Förster resonance energy transfer (FRET) (Kasry et al., 2012). We used a fluorescent cholesterol analog, TopFluor Cholesterol (TFC, a.k.a. cholesterol conjugated to boron-dipyrromethene, BODIPY) and performed fluorescence lifetime imaging microscopy (FLIM). TFC is reportedly very similar to cholesterol in terms of molecular and cellular characteristics (Hölttä-Vuori et al., 2008), and FLIM is a sensitive method for measuring FRET (Bastiaens and Squire, 1999). To provide an unbiased representation of the FLIM data, we applied the phasor approach, which transforms the fluorescence decay at every pixel to a point in a phasor plot (Digman et al., 2008). In the absence of FRET, data points center at a spot on the semicircle, representing single-exponential decays of fluorescence lifetime; when FRET occurs, data points are shifted inside of the semicircle, representing multi-exponential decays (Digman et al., 2008). While 1 μM TFC in 0.02 wt% PVP solution exhibited a single-component lifetime (i.e. >99% points clustered on the semicircle) (Fig. 2b), the addition of GNFs (260 μg/L) dispersed points inside the semicircle (Fig. 2b), suggesting FRET between GNFs and TFC. In contrast, GNFs did not alter BODIPY’s fluorescence lifetime (Fig. 2b), indicating that FRET requires the cholesterol group. We also tested BODIPY-tagged phosphocholine (TFPC) and sphingomyelin (TFSM), the most abundant lipids in the plasma membrane and in nanodomains, respectively. Neither lifetime was significantly affected by GNFs (Fig. 2b and S1), suggesting that GNFs favor cholesterol over PC or SM. To test if TFC outcompetes other lipids in the plasma membrane, we loaded it in live 3T3 cells, a fibroblast cell line widely used for cell membrane studies. After GNFs application, FRET was readily seen in patches across the cell surface and confirmed by phasor analysis (Fig. 2c). GNFs failed to affect membrane-embedded DiO (a common lipophilic fluorescent dye), even after a prolonged incubation (Fig. S2). Taken together, these results suggest that GNFs selectively interact with cholesterol.

**Figure 2.**
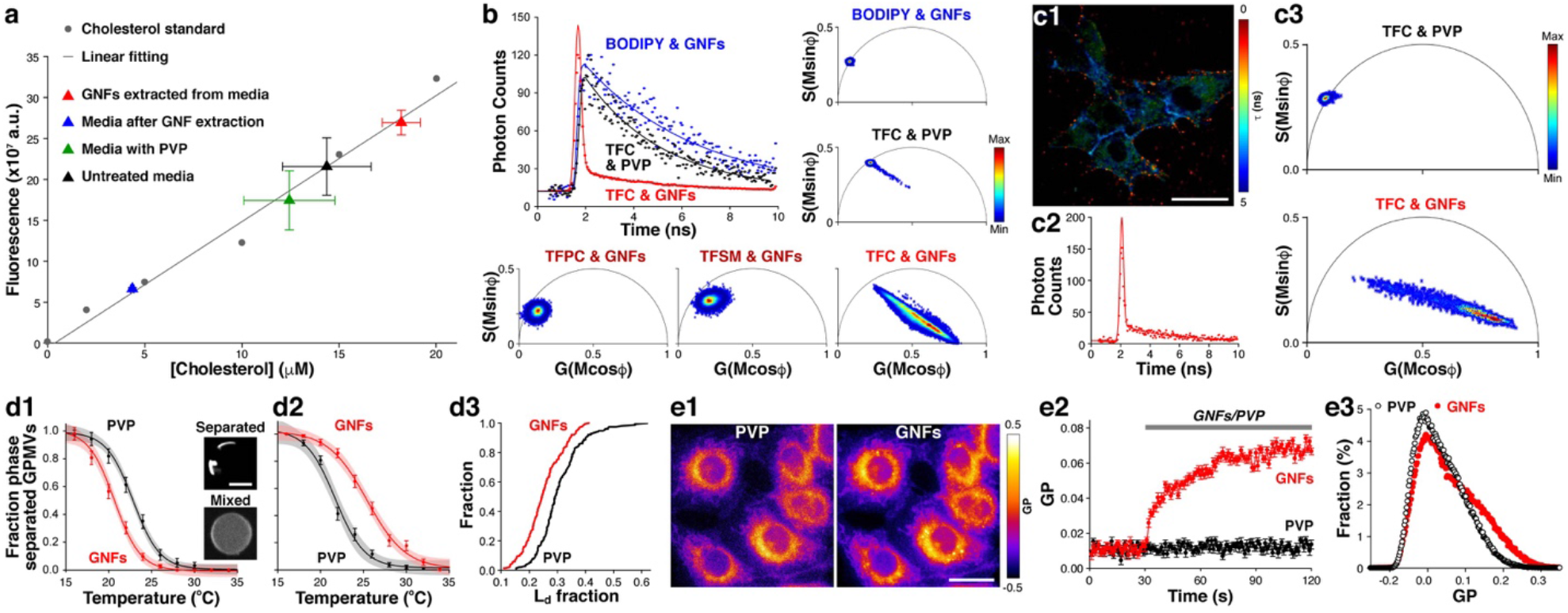
Graphene nanoflakes interact with cholesterol and change cell membrane organization. **(a)** Fluorescent enzymatic assay of cholesterol content in untreated, PVP, or GNF-treated cell culture media. The working curve was generated using cholesterol standards. (b) FLIM data for BODIPY or TFC in the presence of GNFs or PVP controls; best fits are 1 or 2-component exponential decays (in ns, τ_TFC&PVP_ = 3.36, τ_BODIPY&GNFs_ = 4.47, and τ_T_FC&GNFs(1)__ = 0.15 and τ_TFC&GNFs(2)_ = 3.19). FLIM data are also displayed in corresponding phasor plots along with those for GNF-treated TFPC and TFSM. **(c)** FLIM data for TFC loaded in the cell membrane after PVP or GNFs application. **(c1)** Representative pseudo-color shaded FLIM image of TFC-labeled 3T3 cells treated with GNFs. Scale bar, 20 μm. **(c2)** Corresponding FLIM data (red) with a 2-component fit (in ns, τ_1_ = 0.15 and τ_2_ = 3.19). **(c3)** Corresponding phasor plots. **(d)** Temperature-dependent lipid phase separation of GPMVs after PVP (black) or GNFs (red) treatment. **(d1)** T_misc_ of GPMVs isolated from 3T3 cells pretreated with PVP or GNFs for 1 hour. L_d_ is visualized by FAST-DiI. Phase separated fraction was calculated from the numbers of phase-separated and non-separated GPMVs. Area of light shading shows the 95% CI on the sigmoidal fit function. Insets are sample images of phase separated or mixed vesicles. Scale bar, 5 μm. **(d2)** T_misc_ of GPMVs for 1-hour oGNFs or PVP treatment after isolation from 3T3 cells. **(d3)** Cumulative distribution of L_d_ fractions of GPMVs in **(d2)** at 15 °C. **(e)** GP imaging of live 3T3 cells preloaded with C-Laurdan. **(e1)** Sample GP images of same cells treated with PVP (left) and GNFs (right) for 5 minutes each. Scale bar, 10 μm. (e2) Average GP values during PVP (black) and GNFs (red) application. **(e3)** GP distributions in the presence of PVP (open black circles) or GNFs (filled red dots) (*p* < 0.05). Error bars are S.E.M. BODIPY, boron-dipyrromethene; TFC, TopFluor cholesterol (BODIPY-cholesterol); FLIM, fluorescence lifetime microscopy; TFPC, TopFluor phosphocholine; TFSM, TopFluor sphingomyelin; GPMV, giant plasma-membrane vesicle; T_misc_, average miscibility transition temperature; L_d_, disordered liquid phase; GP, generalized polarization.

Few approaches exist to measure cholesterol in the plasma membrane. So to test if GNFs affect membrane cholesterol distribution, we measured their effect on the ability of the plasma membrane to generate co-existing membrane domains, a process which is regulated by cholesterol concentration (Baumgart et al., 2007; Yeagle, 1985). This measure of plasma membrane heterogeneity can be readily monitored in cell-derived giant plasma-membrane vesicles (GPMVs), a preparation with molecular composition and biophysical properties similar to those of the plasma membrane (Sezgin et al., 2012). GPMVs exhibit a temperature-dependent separation into micron-sized lipid ordered (L_o_) and disordered (L_d_) phases (Levental et al., 2009) (Fig. 2d1 insets), and the temperature at which 50% of vesicles contain co-existing L_o_ and L_d_ domains is defined as the average miscibility temperature, T_misc_. We reasoned that pretreatment of cells with GNFs would transiently increase plasma membrane cholesterol levels and thus alter T_misc_. When cells were treated with GNFs for 1 hour before GPMV isolation, T_misc_ of the derived GPMVs was lowered from 27.8 °C in controls to 25.2 °C (Fig. 2d1, Fig. S3), a change similar to increased membrane cholesterol (Levental et al., 2009). Consistently, Filipin staining showed increased surface cholesterol after the 1-hour treatment (Fig. S4). Given that GPMVs, once vesiculated, are isolated from intracellular cholesterol supplies, we carried out a separate experiment to test if GNFs would modulate membrane heterogeneity in isolated GPMVs. Since GNFs can attract cholesterol, we hypothesized that GNFs would preferentially stabilize the cholesterol-rich L_o_ domains (Baumgart et al., 2007; Gray et al., 2013). In line with this prediction, we found that GNFs-treated GPMVs had an increased L_o_ fraction (Fig. 2d3) and an increased T_misc_ (Fig. 2d2 and Fig. S5). Taken together, these data support the idea that membrane-inserted GNFs attract endogenous cholesterol and thus increases the immiscibility of the two phases. To study the biophysical effect of GNFs in live cells, we performed generalized polarization (GP) imaging, which utilizes fluorescent reporters sensitive to membrane lipid order (Barrantes et al., 1999). We labeled 3T3 plasma membranes with C-laurdan, an improved GP probe (Kim et al., 2007), and performed time-lapse imaging. GNFs significantly increased the GP value (Fig. 2e1&3) in a time-dependent manner (Fig. 2e2), suggesting a GNF-induced cholesterol enrichment in the plasma membrane. FLIM, GPMV and GP imaging results collectively demonstrate that GNFs can (1) directly and favorably interact with cholesterol, (2) locally enrich cholesterol in the plasma membrane, and (3) consequently increase membrane lipid packing.

### Graphene Allosterically Modulates Transmembrane Receptors

As membrane cholesterol directly or indirectly regulates many integral proteins (Lingwood and Simons, 2010; Oates and Watts, 2011; Yeagle, 2014), we explored the utility of GNFs in manipulating transmembrane proteins. To do so, we chose ubiquitously expressed and therapeutically important P2Y receptors (P2YRs), a class of GPCRs preferentially localized within cholesterol-enriched nanodomains (N and Volonte, 2013). When activated by extracellular ATP, P2YRs trigger a Ca^2+^-release from internal stores, which can be measured by fluorescent Ca^2+^-indicators using time-lapse imaging (Zhang et al., 2004). In 3T3 cells, we confirmed that P2YRs were the predominant mediator of ATP-induced Ca^2+^-responses (Fig. S6 and supplementary results). Membrane conductance remained unchanged with or without GNFs treatment (Fig. S7), indicating no effect on cell membrane permeability, unlike what was observed in prokaryotes (Tu et al., 2013). Although GP and Filipin imaging suggest that GNFs increase cell surface cholesterol, gas chromatography coupled with a flame ionization detector (GC-FID) mass spectroscopy found no significant changes in total cellular cholesterol, even after a 24-hour treatment (Fig. S8). These results suggest that, instead of changing whole-cell cholesterol metabolism, GNFs enrich cholesterol in the plasma membrane – consistent with the changes in miscibility temperature in the GPMV studies.

As a measure of P2YR activity, we applied two consecutive ATP stimuli at a 1-minute interval, which is insufficient to replenish internal Ca^2+^ stores. Increased P2YR activity should thus make the first response larger but the second smaller. The ratio of 2^nd^ vs. 1^st^ Ca^2+^-responses offers a more reliable readout of P2YR activity, as it is insensitive to the number of surface P2YRs and the capacity of internal Ca^2+^ stores, both of which can be variable among cells. We treated cells for 5 minutes and 1 hour, as these shorter time periods are more likely to result in cell surface localization rather than internalization of GNFs. 5-minute treatment significantly increased the first response and reduced the second (Fig. 3a1), resulting in a smaller 2^nd^ vs. 1^st^ response ratio (Fig. 3a2). 1-hour treatment also led to a smaller ratio, although the 2^nd^ response was slightly larger than that of controls (Fig. 3a). It may be because the 1-hour treatment promotes the refilling of internal Ca^2+^-stores, which is reportedly regulated by membrane cholesterol (Dionisio et al., 2011; Gwozdz et al., 2012). To confirm the involvement of membrane cholesterol in P2YR enhancement, 0.5 mM MβCD was co-applied with GNFs, which diminished the GNF-induced enhancement of Ca^2+^-response (Fig. 3a). In addition, GNFs alone did not cause any Ca^2+^-response (Fig. 3c1), nor did they affect cell membrane permeability (Fig. S7), which further excludes the possibility that GNFs permeabilize the plasma membrane, causing Ca^2+^ influx. To confirm that GNFs’ direct interaction with membrane cholesterol is required, we pre-coated GNFs with 1% W/V SDS (O’Connell et al., 2002), which leads to complete surface coverage of GNFs (Hsieh et al., 2013b) without affecting the dispersion stability (Hsieh et al., 2013a). SDS-coated GNFs failed to increase the ATP-induced Ca^2+^-response, even after 1-hour treatment (Fig. 3b). Furthermore, we either applied ATP and GNFs simultaneously or separately to determine whether GNFs act orthosterically or allosterically. Only co-application of GNFs with ATP potentiated the Ca^2+^-response (Fig. 3c), demonstrating that GNFs behavior as a positive allosteric modulator for P2YRs.

**Figure 3.**
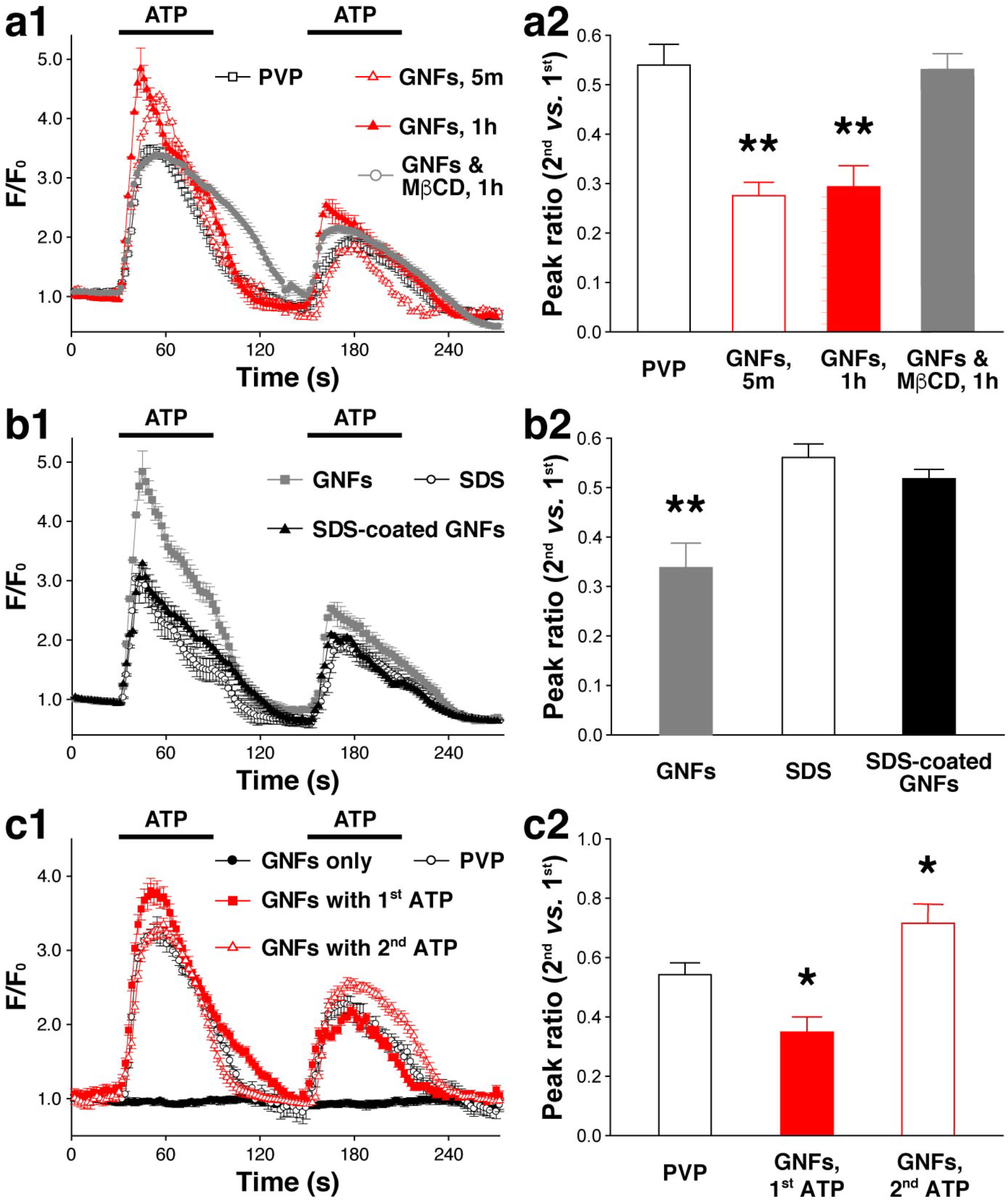
GNFs potentiate P2YR signaling in 3T3 cells. **(a1)** Effect of 5-minute (open red triangles) or 1-hour (filled red triangles) GNFs, PVP (open black squares), or 1-hour GNFs plus MβCD treatment on Ca^2+^-response to 100 μM ATP. **(a2)** Ratios of 2^nd^ to 1^st^ peak amplitude (all n = 6, *p* < 0.01 for 5m and 1h GNFs, and *p* > 0.05 for 1h GNFs & MβCD). **(b1)** Effect of SDS coating of GNFs (black triangles) on ATP-induced Ca^2+^-response, SDS (open circles) and GNFs (gray squares) are shown as controls. (b2) Ratios of 2^nd^ to 1^st^ peak amplitude (all n = 6, and *p* > 0.05 for SDS&GNFs vs. SDS, *p* < 0.01 for SDS&GNFs vs. GNFs). **(c1)** Co-application of GNFs with either the 1^st^ (filled red squares) or 2^nd^ (open red triangle) ATP application, GNFs alone (filled black circles) and PVP (open black circles). **(c2)** Ratios of 2^nd^ to 1^st^ peak amplitude (all n = 6, *p* < 0.05 for both GNFs co-applications vs. PVP). \**p* < 0.05, \*\**p* < 0.01. MβCD, methyl-β-cyclodextrin, ATP, adenosine triphosphate, SDS, sodium dodecyl sulfate.

### Graphene Promotes Neurotransmitter Release

In addition to the modulation of membrane proteins, cholesterol is also critical to the secretion of signaling molecules. One of the best examples is the highly choreographed release of neurotransmitter from synaptic vesicles in nerve terminals. Emerging evidence has shown that cholesterol is essential for the origination and maintenance of synaptic vesicle pools (Dason et al., 2014; Mauch et al., 2001), for the localization and function of exo- and endocytotic proteins, and for the fluidity and curvature of vesicular and plasma membranes (Chang et al., 2009; Kreutzberger et al., 2015). Therefore, we asked if GNFs could be used to modulate neurotransmitter release. Here we used hippocampal cultures, an established preparation for studying neurotransmission. First, we tested if GNFs could elevate cholesterol within neuronal membranes, which already contain high levels of cholesterol. In nearly all treated neurons, we observed a significant increase of GP value after 1 hour GNF treatment (Fig. 4a1), although the increase was slower and smaller than that seen in astrocytes (Fig. 4a2). As a gross measure of functional impact, we performed whole-cell patch clamp recordings and measured miniature evoked postsynaptic currents (mEPSCs) as well as postsynaptic NMDA- and AMPA-receptor mediated currents (I_NMDAR_ and I_AMPAR_). Between GNF-treated neurons and controls, there was no significant difference in mEPSC amplitude or I_NMDAR_ vs. I_AMPAR_ ratio (Fig. S9). Inter-event interval, however, was significantly reduced after GNF treatment (Fig. S9a). Taken together, this indicated that the effect of GNFs was presynaptic rather than postsynaptic.

**Figure 4.**
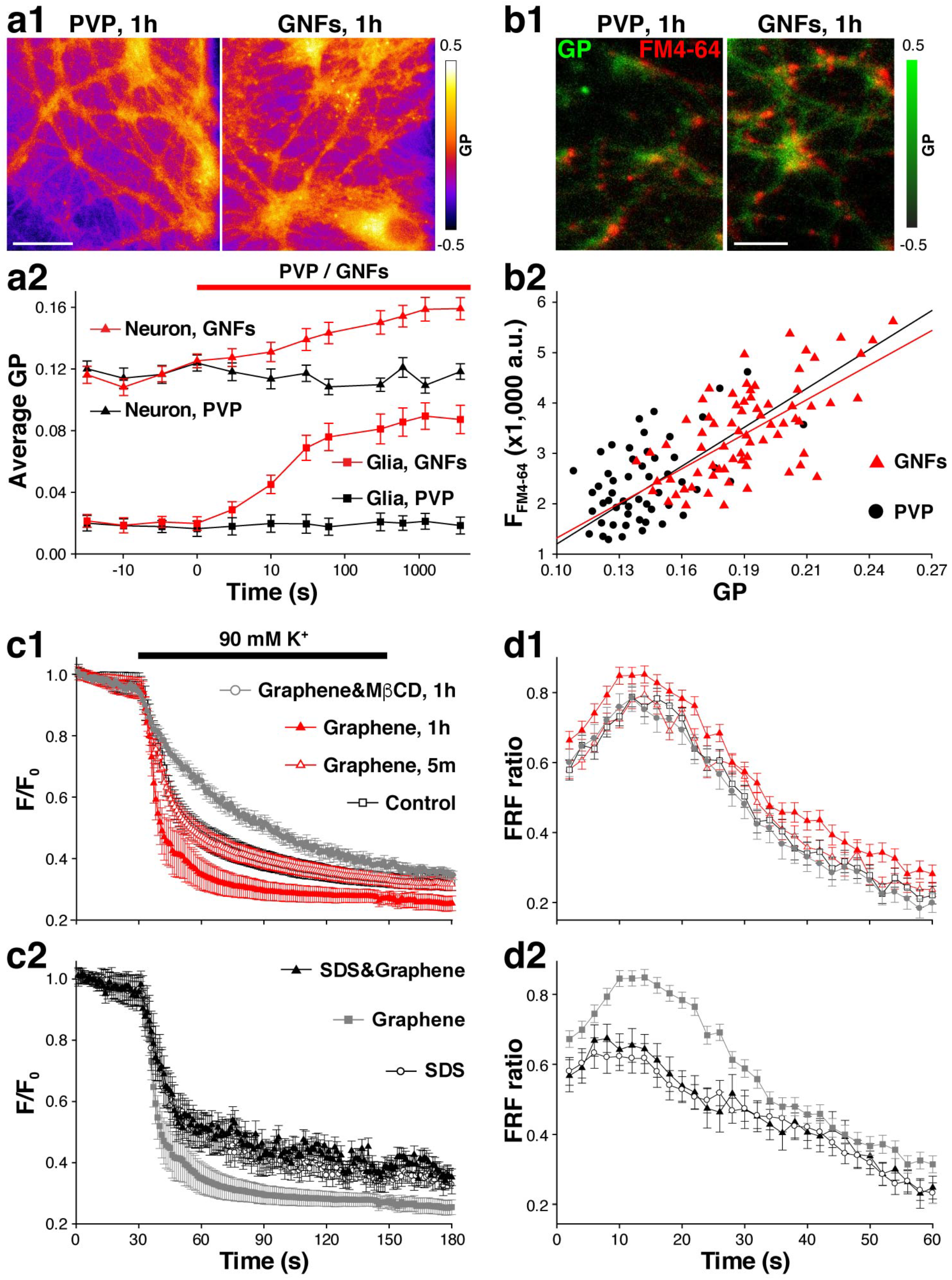
GNFs potentiate neurotransmission by increasing membrane cholesterol. **(a)** GP imaging of neurons. **(a1)** Sample GP images of neurons after 1 h treatment with either PVP (left) or GNFs (right). Scale bar, 20 μm. **(a2)** Time-correlated GP value for PVP (black) or GNFs (red) application for both neurons (triangles) and glia (squares). **(b)** C-Laurdan and FM4-64 colabelling. **(b1)** Sample GP (green) images of neurons after 1 h treatment with either PVP (left) or GNFs (right) are superimposed with FM4-64 (red) images of the same fields of view. **(b2)** Scatter plot of FM4-64 intensity vs. GP value for GNFs (red triangles) and PVP (black circles). Linear regression fits have R of 8.821 for GNFs and 7.924 for PVP. **(c)** FM destaining with 2 minutes 90 mM K+. **(c1)** The destaining after 5-minute (open red triangles) or 1-hour (filled red triangles) treatment with GNFs or 1-hour GNFs & MβCD (open black circles) co-treatment are compared with 1-hour PVP treatment (open black squares) (all n = 6; for 5 min, *p* > 0.05; for 1 h, *p* < 0.05; for GNFs & MβCD, *p* > 0.05). (c2) Destaining after 1-hour (filled gray squares) treatment with GNFs is significantly faster than 1-hour SDS (open circle) or SDS-coated GNFs treatment (filled black triangles) (all n = 6; both *p* < 0.05). **(d)** Qdot-based measurement of fusion modes during 1 min 10 Hz electric field stimulation, shown as the ratio of FRF vs all fusion events. **(d1)** Four conditions same as **(c1)** with the same symbol sets (all n = 6; between PVP and 5-minute GNFs, *p* > 0.05; between PVP and 1-hour GNFs, *p* < 0.05; between PVP and GNFs & MβCD, *p* > 0.05). **(d2)** Same conditions as **(c2)** with the same symbol sets (all n = 6; both*p* < 0.05). Qdot, quantum dot; FCF, full-collapse fusion; FRF, fast and reversible fusion.

To directly examine if there were presynaptic changes, we used FM4-64, whose loading and unloading reflect the amount of releasable vesicles and their release probability (P_r,v_), respectively (Betz and Bewick, 1992). 1-hour GNFs treatment increased FM4-64 uptake (Fig. 4b1), suggesting an enhancement of activity-evoked vesicle retrieval. FM4-64 fluorescence intensity was correlated to GP value at synaptic boutons (Fig. 4b2), suggesting that the GNFs’ presynaptic effect is associated with increased membrane cholesterol. During subsequent field stimulation, the rate of FM4-64 destaining in GNF-treated neurons was significantly faster (Fig. 4c1), indicating a higher release probability (P_r,v_) and/or a larger fraction of releasable synaptic vesicles. Again, pre-coating of GNFs with SDS abolished these effects (Fig. 4c2), demonstrating that a direct GNF-cholesterol interaction is required.

To further illustrate the changes in synaptic vesicle turnover, we turned to quantum dot (Qdot) enabled single vesicle imaging (Zhang et al., 2009). The hydrodynamic diameter of the Qdots we used was ~15 nm, smaller than the luminal diameter of a synaptic vesicle (~25 nm) but much larger than the estimated fusion pore size (~1-3 nm). This restricted the loading to one Qdot per synaptic vesicle (Zhang et al., 2007), which allowed for an accurate estimate of the total releasable pool (TRP) of vesicles at each synapse. We calculated the total number of Qdots taken up after maximal loading (Fig. S10a) – i.e. strong stimulation (2 minutes, 90 mM K+) with a high concentration of Qdots (100 nM) (Zhang et al., 2009). In good agreement with FM4-64 data, 1-hour treatment, on average, resulted in a 12% increase of releasable synaptic vesicles (Fig. S10b). This is also consistent with the time needed to achieve a significant increase of neuronal GP value. Using time-lapse single Qdot imaging, we analyzed the turnover kinetics of synaptic vesicles. We applied the same stimulation but used a much lower concentration of Qdots (0.8 nM), which resulted in the random labeling of individual vesicles across the TRP (Zhang et al., 2009). Quantal analysis of Qdot photoluminescence in FM4-64 defined individual synaptic boutons (Zhang et al., 2009) confirmed single vesicle loading (Fig. S10c). Neurons were perfused for 15 minutes to remove excess Qdots and a 10-Hz 60-s electric field stimulation was applied to evoke synaptic vesicle release. Deconvolution of Qdot photoluminescence changes allowed us to distinguish between two different modes of vesicle turnover, the classical full-collapse fusion (FCF), or fast and reversible fusion (FRF): a small increase immediately followed by the complete loss of unitary Qdot photoluminescence or a small increase alone, respectively (Zhang et al., 2009) (Fig. S10d). After 1-hour GNF treatment, there were significantly more FRF events during stimulation and an increase in the number of FRF events per vesicle (Fig. S10f), leading to a higher FRF ratio (Fig. 4d1). Furthermore, the time to the first fusion event for individual vesicles was significantly shorter in GNF-treated neurons (Fig. S10e), in good agreement with a higher P_r,v_. To confirm that these changes in synaptic vesicles were indeed due to GNF-induced cholesterol enrichment, we co-applied MβCD, which offset the GNF-induced changes in synaptic vesicle turnover as measured by both FM and Qdot imaging (Fig. 4c&d). To verify the necessity of a direct interaction between GNFs and plasma membrane cholesterol, we again used SDS-coated GNFs. As expected, pre-coated GNFs failed to induce vesicular changes (Fig. 4c2 and d2). Altogether, these results suggest that, by inducing local cholesterol enrichment, GNFs can promote neurotransmission in a presynaptic fashion, illustrating a new method to enhance neurotransmission.

## Discussion

Recent advances in single molecule tracking, protein engineering, and super-resolution imaging have provided the ability to monitor nanodomain dynamics at previously unprecedented spatiotemporal resolution (Simons and Gerl, 2010). However, progress in manipulating nanodomains with matching precision has been far more limited. Particularly, the ability to stabilize the L_o_ phase in the plasma membrane of live cells would provide valuable insight into the existence, organization, and functionality of membrane nanodomains(Lingwood and Simons, 2010). Here, we provide independent lines of evidence to demonstrate a direct interaction between GNFs and cholesterol. GNFs attract membrane cholesterol upon insertion into cell membranes, which leads to the formation or stabilization of cholesterol-enriched L_o_ domains and thus changes the architecture and biophysical properties of the plasma membrane. Unlike pharmacological and genetic tools, such as statins or the NPC1 mutation, GNFs act on cell membranes directly and acutely, rearranging endogenous membrane cholesterol with little impact on total cell cholesterol. In contrast to existing biochemical tools like cyclodextrins, GNFs offer a faster and more targeted way to enrich cholesterol in the plasma membrane. In different cell types and for different signaling pathways, the effect of GNFs is determined by the specific roles membrane cholesterol plays.

Here, we have used two different cell preparations to demonstrate the utility of GNFs. For signal transduction via transmembrane receptors like P2YRs, GNFs can allosterically promote their activity, likely because of the potentiating role cholesterol plays in receptor localization, trafficking, stability, and dimerization (Cherezov et al., 2007; Hanson et al., 2008; N and Volonte, 2013). The time needed for GNFs to enhance P2YRs matches that of the GNF-induced GP increase in live cells (i.e. both in seconds), suggesting that lipid phase stabilization contributes to the receptor enhancement ^42^. Equally possible is that elevated cholesterol promotes GPCR activity via direct binding, which facilitates receptor dimerization or association with downstream factors (Cherezov et al., 2007; Hanson et al., 2008). For the secretion of signaling factors, such as neurotransmitters in the nervous system, GNFs can potentiate release by promoting vesicle origination, recycling, and membrane fusion/fission, all of which are potentiated by increased membrane cholesterol (Chang et al., 2009; Dason et al., 2014; Kreutzberger et al., 2015; Mauch et al., 2001). While reducing cholesterol in neurons with conventional tools like MβCD demonstrated its importance to synaptic plasticity via modulation of ion channels ^61^ and receptors ^62^, GNFs, by increasing synaptic membrane cholesterol, reveal new routes to achieve presynaptic potentiation (i.e. by accelerating synaptic vesicle turnover).

We observed a difference between 3T3 cells and neurons in the time courses of GNF’s effects, possibly due to a significant difference of initial membrane cholesterol concentration. In support of this idea, astrocytes, similar to 3T3 cells (with less membrane cholesterol than neurons), exhibited a significantly faster and larger average increase of GP value than neurons (Fig. 4a2). It is also possible that the stochastic and diffusion-limited process of unmodified GNFs reaching synapses underlies their slower presynaptic effects. As better surface functionalization and conjugation strategies for carbon nanomaterials are rapidly evolving (Mao et al., 2013), the conjugation of GNFs with biomolecules like antibodies or ligands may allow the rapid and selective targeting of GNFs to specific subcellular sites, achieving a spatiotemporal precision suitable for the manipulation of highly dynamic membrane nanodomains or even the membranes of subcellular organelles. Furthermore, graphene-based cholesterol manipulation will accelerate the investigation of cholesterol-related disorders. In particular, site-specific modification of cholesterol will be informative in understanding the pathological contribution of membrane cholesterol in related diseases (Maxfield and Tabas, 2005), such as Niemann Pick type C disease and Alzheimer’s disease.

## Author Contributions

K.E.K. and Q.Z. designed experiments with input from all authors and wrote the manuscript. K.E.K. and Q.Z. performed all imaging and spectrofluorometry experiments. R.M.L. performed all electrophysiology experiments. K.E.K and J.R.M. performed whole-cell cholesterol quantification. K.E.K. and A.T.S. performed all FLIM measurements with input from M.C.S. and Q.Z. K.E.K. and K.R. performed all GPMV experiments with input from A.K.K. and Q.Z. T.H., Y.Z. and Y.Q.X. prepared GNFs, and performed all TEM, Raman spectroscopy and AFM measurements. J.A.M. supervised cholesterol quantification, A.K.K. supervised GPMV experiments, M.C.S. supervised FLIM experiments, and Q.Z. supervised all experiments.

## Acknowledgment

We thank I. Kristaponyte for technical assistance with cell culture. We thank Joseph T. Sharick for assistance in FLIM data acquisition and analysis. We thank H.E. Hamm, A. H. Brown, and K.P.M. Currie for valuable comments and discussions. We thank all members of the Zhang and Xu laboratories for their support.

## Abbreviations

GNF: graphene nanoflake;
PVP: polyvinylpyrrolidone;
TEM: transmission electron microscopy;
AFM: atomic force microscopy;
BODIPY: boron-dipyrromethene;
TFC: TopFluor cholesterol (BODIPY-cholesterol);
FLIM: fluorescence lifetime microscopy;
TFPC: TopFluor phosphocholine;
TFSM: TopFluor sphingomyelin;
GPMV: giant plasma-membrane vesicle;
T_misc_: average miscibility transition temperature;
L_d_: disordered liquid phase;
GP: generalized polarization;
MβCD: methyl-β-cyclodextrin,
ATP: adenosine triphosphate,
SDS: sodium dodecyl sulfate;
Qdot: quantum dot;
FCF: full-collapse fusion;
FRF: fast and reversible fusion.

## Supplementary Materials

### Materials and Methods

#### Preparation and Characterization of Suspended Graphene Nanoflake (GNF) Solutions

The GNF suspension was prepared by liquid exfoliation of graphite powder. Graphite powder was purchased from ASBURY CARBONS (Grade: 2299). Polyvinylpyrrolidone (PVP, MW: 1,300,000 g/mol) was purchased from Sigma. Hydrophobic graphite powder was added into 2 wt% PVP or 1 wt% SDS (sodium dodecyl sulfate) water solution and sonicated for 9 h in a bath sonicator^1^ The uniform GNF suspension was then centrifuged with a Thermo Scientific Fiberlite F15-6 X 100y rotor at 4,000 rpm and at room temperature for 1 h to sediment large graphite aggregates. The upper 50% of supernatant was carefully decanted, resulting in PVP-functionalized GNF suspension^2,3^. The transmission characterization of GNF suspension was carried out on a Varian Cary 5000 UV-VIS-NIR spectrophotometer. The concentration of GNF was estimated with an absorption coefficient of 2460 L g^−1^ m^−1^ at 660 nm ^2^, which is typically 26 mg/L for freshly-made GNF suspension. The suspension is stored in 4 °C and remain stable for more than a year. As shown in Figure 1, a one-year old GNF suspension *(left)* shows no precipitation and evenly-distributed as that of a freshly-prepared one *(right).*

#### Characterization of GNFs

The GNF suspension was characterized using transmission electron microscopy (TEM). TEM samples were prepared by drop casting a small volume (~ 2 μl) of GNF suspension onto carbon grids (300 mesh size, Ted Pella). The samples were air dried for 2 h, and then rinsed with DI water to remove excessive solvent. Bright field TEM images of the typically observed GNFs were taken by an Osiris TEM at an accelerating voltage of 200 kV. Raman spectroscopy was also employed to characterize the quality of the GNFs. The GNFs were drop-casted onto an Si/SiO2 wafer (300 nm SiO_2_) and then washed with DI water to remove excess suspension agents. The Raman spectra were taken by a DXR Raman microscope (Thermo Scientific) with 532 nm laser excitation. The size and thickness of GNFs were further investigated by a Nanoscope III atomic force microscope (AFM). The GNF suspension was spin-coated onto a Si wafer with 300 nm SiO_2_, and washed with DI water to remove solvent residue. The AFM was operated in tapping mode with a typical image size of 2-5 μm.

#### Cell Culture

For all experiments, rat postnatal hippocampal cultures were prepared as previously described ^4^, with some modifications. Briefly, rat hippocampi (CA1-CA3) were dissected from P0 or P1 Sprague-Dawley rats and dissociated into a single-cell suspension with a 10 min incubation in Trypsin-EDTA (Life Technologies) followed by gentle trituration using three glass pipettes of different diameters (~ 1 mm, 0.5 mm, and 0.2 mm), sequentially. Dissociated cells were recovered by centrifugation (x 200 g, 5 minutes) at 4 °C and re-suspended in plating media composed of Minimal Essential Medium (MEM, Life Technologies) with (in mM) 27 glucose, 2.4 NaHCO_3_, 0.00125 transferrin, 2 L-glutamine, 0.0043 insulin and 10%/vol fetal bovine serum (FBS, Omega). 100 μl of cell suspension was added onto round 12mm-∅ glass coverslips (200-300 cells/mm^2^) 100 μl of Matrigel (BD Biosciences, 1:50 dilution) was deposited on the coverslips and incubated at 37°C with 5% CO_2_ for ~ 2 h, then aspirated before cells were plated. Cells were allowed to settle on the coverslip surfaces for 4 h before the addition of 1 mL culture media made of MEM containing (in mM) 27 glucose, 2.4 NaHCO3, 0.00125 transferrin, 1.25 L-glutamine, 0.0022 insulin, 1 %/vol B27 supplement (Life Technologies) and 7.5 %/vol FBS. 1 to 2 days after plating, 2% Ara-C was introduced with another 1 mL of culture media, which efficiently prevented astroglia proliferation. All procedures were in accordance with Vanderbilt University Institutional Animal Care and Use Committee standards.

NIH 3T3 cells were grown at 37°C with 5% CO_2_ in Dulbecco’s modified Eagle’s medium containing 4.5 g/L glucose and L-glutamine supplemented with 10% fetal bovine serum, 100 units mL^−1^ penicillin, and 100 μg mL^−1^ streptomycin. Cells were regularly passaged to maintain adequate growth and were passaged at least 5 times before trypsinization and plating on the Matrigel-coated round 12mm-∅ glass coverslips (75 μL of 1-3 × 10^6^ cell solution per coverslip). Cells were grown to 50-80% confluency for 24 h on coverslips prior to experiments.

#### Cholesterol Assay

An enzymatic assay (Amplex Red Cholesterol Assay Kit, Life Technologies) was used to quantify free cholesterol concentrations^5^. Briefly, a serial dilution of cholesterol standard was used to generate a calibration curve. Four different batches of hippocampal culture media were used. Aliquots of media were incubated with H_2_O, 0.002 wt% PVP in H_2_O, or 260 ng/mL GNFs with 0.002 wt% PVP in H_2_O in 37°C and 5% CO_2_ for 24 h. Graphene was separated from media using Amicon centrifugal filters (30 kDa, Millipore) and resuspended in H_2_O with 0.002 wt% PVP. The flow-through media was collected. Media with H_2_O, media with PVP, or GNFs separated from media and flow-through media were assayed, and the fluorescence intensities of all enzymatic reaction products were measured using a spectofluorometer (FluoroMax-4, Horiba). Four independent measurements were performed for each sample and the cholesterol concentrations were calculated based on the calibration curve.

#### Fluorescence lifetime measurements

Fluorescence lifetime images were acquired using a custom-built multiphoton fluorescence system (Bruker) built on an inverted microscope (Nikon Ti-E). For bulk solution measurements, 500 μL sample volumes were illuminated using a 40x oil-immersion objective (N.A. 1.3). TFC (TopFluor Cholesterol) or BODIPY (boron-dipyrromethene) were added to suspended GNFs (26 ng/mL) or water at a final concentration of 1 μM for 1 h prior to imaging. NIH-3T3 cell samples were illuminated using a 100x oil-immersion objective (N.A. 1. 45). For cell membrane experiments, graphene suspension (26 ng/ml) was added to cells in normal Tyrode for 10 min and then TFC was added at a final concentration of 1 μM. Samples were excited with a Ti:Sapphire laser (Coherent, Inc.) tuned to 960 nm, passed through a 550/100 emission filter, and detected using a GaAsP photomultiplier tube (H7422P-40, Hamamatsu). Pixel dwell time was 4.8 μs and the acquired images were 256×256 pixels with a 60 s acquisition time at an average incident power of approximately 10 mW. Fluorescence lifetime images were acquired using time-correlated single photon counting electronics (SPC-150; Becker & Hickl). The instrument response function (IRF) was generated by measuring the second harmonic generation of urea crystals excited at 900 nm. The full width at half maximum of the IRF was 244 ps. Fluorescence lifetime was validated before each experiment by imaging a fluorescent bead standard (Polysciences, Inc.). The measured lifetime of this bead was 2.1 ± 0.005 ns (n=3), in good agreement with published values ^6^. Phasor analysis was performed as described previously ^7^, using Matlab algorithms based on a previously published algorithm ^8^.

#### Giant plasma membrane vesicle preparation

NIH-3T3 cells at greater than 85% confluency were washed 3 times with PBS and labeled at 5 ug/ml for 10 min at 37°C with 1,1’-Dilinoleyl-3,3,3’,3’-tetramethylindocarbocyanine,4-chlorobenzenesulfonate (FAST-DiI, C-18, Life Technologies). GPMV isolation was performed as previously described^9^. Briefly, cells were blebbed in deionized water containing, in mM: 10 HEPES, 150 NaCl, 2 CaCl_2_, pH 7.4, with 25 mM PFA and 2mM DTT at 37°C for 2-3 h. GPMVs were treated for either prior to or following vesiculation. For treatments prior to GPMV isolation, the supernatant was collected and allowed to settle for at least 1 hour at 4°C prior to imaging to allow GPMVs to settle. Three independent GPMV preparations were performed for each treatment duration and a minimum of 60 GPMVs were imaged for each individual temperature point.

#### GPMV fluorescence imaging and lipid phase separation

Lipid phase separation was achieved using a commercial liquid-nitrogen cooled temperature stage equipped with two heating elements (GS350, Linkam Scientific Instruments). Thermal grease (Linkam) was applied to stage surfaces for each sample to minimize thermal contact resistance and maximize conductive heat transfer. Samples were allowed to equilibrate for 1 min at each temperature point prior to image acquisition. Images were acquired with a 60X LUMPPlanFl Olympus water-immersion objective (N.A. 0.9) on a customized spinning disk confocal setup built on an Olympus BX-51WI microscope with a CSU-X1 (Yokogawa) spinning disk head and an Evolve 512 EMCCD (Photometrics). Samples were visualized with a 561 nm laser (Coherent) and an Em 605/52 filter set. Image analysis was performed in Matlab using custom-written algorithms.

#### Generalized polarization image analysis

Similar to previous report^10^, cells were pre-incubated with culture media containing 1 μM C-laurdan (TP Probes) at 37°C and 5% CO_2_ for 1 h. Loaded cells were imaged using a Nikon Ti-E equipped with an iX897 EMCCD (Andor) and a 100X ApoVC objective (N.A. 1.40). Cells on coverslips were mounted in an RC-26G imaging chamber (Warner Instruments) bottom-sealed with a 24X40 mm size 0 cover glass (Fisher Scientific). The chamber was fixed in a PH-1 platform (Warner Instruments) placed on the microscope stage. The temperatures of both the imaging chamber and the constant perfusion (~50 μL/sec) solution were maintained at 34°C by a temperature controller (TC344B, Warner Instruments). A filter combination of D350x for excitation and a DiC 409LP with an Em 440/40 or 483/32 (for blue or green channels, respectively) were used. Image acquisition and synchronized perfusion were controlled via Micro-manager software. Images were post-processed in ImageJ using the following formula: I_GP_=(I_blue_-G×I_green_)/(I_blue_+G×I_green_), in which G is the sensitivity correction factor between the two channels^11^. G was empirically determined by imaging 1 μM C-laurdan diluted in DMSO using the standard protocol. Given GP_DMSO_ = 0.006, the G value of our imaging setup was calculated using the following formula: G=(I_blue_×(1-GP_DMSO_))/(I_green_×(1+GP_DMSO_)).

#### Folch Extraction

500 μL and 250 μL of 5% HCl were added to cells and to 250 μL of media, respectively. 750 μL of Folch solution (2:1, CHCl_3_:MeOH with 17 mg/L BHT, butylated hydroxytoluene) was utilized for extraction and 10 μL of 1.25 mg/mL 5β-cholestan-3α-ol was added as an internal standard for cholesterol quantitation. The Folch solution was vortexed and centrifuged briefly to allow distinct organic and aqueous layers to separate. The organic layer was then used for cholesterol identification and quantification (GC-FID, Gas chromatography – Flame ionization detector, and GC-MS, Gas chromatography – Mass spectrometry).

#### Cholesterol derivatization

Folch extractions from both cells and media were dried down and reconstituted in 40 μL of bistrimethylsilyltrifluoroacetamide (BSTFA) kit solution (Sigma-Aldrich) for at least 2 h with internal standard to account for extraction and derivatization efficiencies.

#### Gas chromatography – Flame ionization detector (GC-FID)

Cholesterol quantification was carried out in duplicates for all biological replicates with a GC-6890 gas chromatograph (Hewlett-Packard) equipped with a DB-5 (30 mm × 0.32 mm × 0.25 mm) fused silica column (Sigma Aldrich). Briefly, sterols were separated using a temperature program as follows: samples were heated from 220 to 275 °C at 15 °C/min, then further heated to 280 °C at 1 °C/min and maintained for 2 min, followed by heating to 290 °C at 5 °C/min rate and holding for 10 min.

#### Determination of total protein content

Proteins separated during Folch extraction were MeOH washed, pelleted, and dried before re-suspension in 2% SDS (Sigma Aldrich). A Pierce BCA assay (Life Technologies) was performed according to manufacturer specifications using 25 μL of protein sample per microwell. Sample absorption was measured at 560 nm using a Glomax Discover (Promega) 96-well microplate reader.

#### Filipin Staining

Cells were fixed in PBS containing 4% paraformaldehyde for 1 h, washed with PBS, and incubated with filipin (1:500 in PBS, Sigma-Aldrich). For whole cell staining, cells were incubated with filipin for 2 h at room temperature. For membrane staining, cells were incubated with filipin for 30 min at 4 °C to minimize membrane penetration. Fluorescence imaging was performed on an Olympus IX-81 inverted microscope using an Olympus PlanSApo 20X objective (N.A. 0.75) with a fluorescence filter set (Ex 390/40, DiC T425LPXR, Em 460/50, Semrock). Images were acquired with an EMCCD (Andor) via Micro-manager with the same acquisition settings (excitation light intensity, exposure time and gain) across all samples. For analysis, three independent batches of cultures and at least three cells for every group were analyzed (n > 9 different coverslips). Image analysis was performed in both ImageJ and Matlab using custom-written algorithms.

#### Fluorescence imaging and analysis

All live cell imaging was performed using the spinning disk confocal setup used for GPMV imaging. Cells on coverslips were mounted in an RC-26G imaging chamber (Warner Instruments) bottom-sealed with a 24X40 mm size 0 cover glass (Fisher Scientific). The chamber was fixed in a PH-1 platform (Warner Instruments) placed on the microscope stage. Gravity perfusion was controlled by a VC-6 valve control system (Warner Instruments) with a constant rate of ~50 μL/sec. All perfusion lines were combined into an SHM-6 in-line solution heater (Warner Instruments). The temperatures of both the imaging chamber and the perfusion solution were maintained at 34°C by a temperature controller (TC344B, Warner Instruments). Image acquisition and synchronized perfusion were controlled via Micro-manager software.

For TFC loading, cells were pre-incubated in culture media containing 1 μM TFC at 37° C with 5% CO_2_ for 20 min. For DiO imaging, cells were pre-loaded with 10 μM DiO (Invitrogen) in culture media at 37°C with 5% CO_2_ for 15 min. For voltage imaging, the DiO containing solution was replaced with 20 mM DPA (dipicrylamine) in normal Tyrode. For Ca^2+^ imaging, cells were pre-incubated with culture media containing 10 μM XRhod-1AM (Life Technologies) at 37°C with 5% CO_2_ for 30 min. For FM dye or Quantum dot (Qdot) loading of the evoked pool of synaptic vesicles, cells were incubated with 10 μM FM1-43 or FM4-64, or 100 or 0.8 nM Qdots (Qdot 605, Life Technologies) for 2 min in high K+ bath solution containing (in mM): 64 NaCl, 90 KCl, 2 MgCl_2_, 2 CaCl_2_, 10 N-2 hydroxyethyl piperazine-n-2 ethanesulphonic acid (HEPES), 10 glucose, 1 μM TTX, pH 7.35. After loading, cells were washed with normal bath solution containing 10 μM NBQX (2,3-dihydroxy-6-nitro-7-sulfamoylbenzo[f]quinoxaline-2,3-dione) and 20 μM D-AP5 (D-(-)-2-Amino-5-phosphonopentanoic acid) for at least 10 min prior to imaging. For TFC, FM1-43 and DiO/DPA imaging, a 480 nm laser (Coherent) and a filter combination of DiC 500LX and Em 520/20 were used. For Ca^2+^ imaging, a 561 nm laser (Coherent) and a filter combination of DiC 580LPXR and Em 605/52 were used. For Qdot imaging, a 480 nm laser and a filter combination of DiC 510LX and Em 605/10 were used. For FM4-64 imaging, a 561 nm laser (Coherent) and a filter combination of DiC 600LX and Em 620/20 were used. All optical filters and dichroic mirrors were purchased from Chroma or Semrock. The acquisition rate was 5 Hz for Qdots and 1 Hz for all other dyes. For each dye, images were taken with the same acquisition settings (excitation light intensity, spinning disk speed, exposure time, and EM gain) for all samples. All image analyses were performed in ImageJ as described previously ^12^. To obtain mean Qdot photoluminescence intensities in each synaptic bouton across all fields of view, all ROIs defined by retrospective FM4-64 staining of the same field of view were projected to an average image made from the first ten frames of the Qdot image stack. Quantal analysis for single Qdot loading was applied as described before ^12^. To analyze the behavior of vesicles labeled by single Qdots, we selected ROIs having only one Qdot. Time-dependent Qdot photoluminescence changes were extracted with a 5-frame moving window. All data were exported and processed in Excel, Matlab, and SigmaPlot.

#### Electrophysiology

Whole-cell voltage clamp recordings were performed on neurons from 12 – 18 DIV cultures using a Multi-Clamp 700B amplifier, digitized through a Digidata 1440A, and interfaced via pCLAMP 10 software (all from Molecular Devices). All recordings were performed at room temperature. Cells were voltage clamped at −70 mV for all experiments. Patch pipettes were pulled from borosilicate glass capillaries with resistances ranging from 3 – 6 MΩ when filled with pipette solution. The bath solution (Tyrode’s saline) contained (in mM): 150 NaCl, 4 KCl, 2 MgCl_2_, 2 CaCl_2_, 10 HEPES, 10 glucose, pH 7.35. The pipette solution contained (in mM): 120 Cesium Methanesulfonate, 8 CsCl, 1 MgCl_2_, 10 HEPES, 0.4 EGTA, 2 MgATP, 0.3 GTP-Tris, 10 phosphocreatine, QX-314 (50 μM), 5 biocytin (Tocris), pH 7.2. For the recordings of mEPSCs, bath solution was supplied with 1 μM tetrodotoxin (TTX, Abcam). The last 50 mEPSCs at the end of 5 min recordings with TTX were collected and analyzed using a template based event detection feature of Clampfit 10.2 software. The template was generated by averaging a collection of representative mEPSC events. To measure AMPA receptor currents, D-AP5 (Abcam), an NMDA receptor antagonist, was added to the bath solution. NMDA receptor currents were recorded in the presence of 10 μM NBQX (Abcam), an AMPA receptor antagonist, in 0 mM [Mg^2+^] / 3 mM [Ca^2+^] bath solution at −70 mV holding potential. Isolated AMPA and NMDA EPSCs were recorded from the same neurons sequentially by first applying D-AP5 then completely replacing it with NBQX. The INMDAR/IAMPAR ratio for every neuron was calculated from the average amplitudes of the last 10 NMDA and AMPA events during 5 min D-AP5 or subsequent NBQX application. No postsynaptic currents were detected if D-AP5 and NBQX were applied together. All signals were digitized at 20 kHz, filtered at 2 kHz, and analyzed offline with Clampfit software (Molecular Devices). All data were exported to and processed in Microsoft Excel.

#### Data Analysis

All experiments were carried out blindly and repeated in at least three different batches. All values presented are mean ± s.e.m. For calculating statistics, the Student’s t-test was used for 2-group comparison of average values, and one-way analysis of variance (ANOVA) and the Tukey-Kramer method as *post-hoc* analysis were used for 3 or more groups. Fisher z-tests were used to compare correlation coefficients. 2-sided *Kolmogorov-Smirnov* tests were used to test for equivalent distributions.

### Supplementary Results

#### Graphene nanoflakes have a limited impact on the electric properties of the cell membrane

Given the high charge carrier mobility of GNFs, they may change the electrical properties of the plasma membrane (e.g. conductance and excitability) when it attaches to or inserts. However, our results exclude this possibility. First, recordings of neuronal mEPSCs in synaptically matured neurons showed no significant difference in their amplitude between the GNF-treated and PVP-treated groups (Fig.S9). Second, we monitored neuronal membrane potential changes using an optical measurement, in which dipicrylamine (DPA, a small lipophilic anion) quenches membrane-embedded DiO upon depolarization ^13^. The presence of GNFs did not cause any significant changes in 90 mM K+-induced neuronal depolarization (Fig. S2b), suggesting that GNFs did not affect neuronal membrane conductance or excitability. Third, we recorded I-V curves of 3T3 cells in the presence of GNFs or PVP control. There was no significant difference between the GNFs and PVP control groups in terms of membrane conductance (Fig. S7), suggesting that GNFs neither breaks cell membrane nor changes ion conductance of the plasma membrane. Together, these data suggest that, although it may insert into the plamsa membrane, GNFs does not compromise membrane integrity or its electrophysiological properties.

#### P2Y receptors mediate ATP-triggered intracellular release of Ca^2+^ from internal stores

Purinergic receptors, including P1 receptors for adenosine and P2 receptors for ATP, are a large family of transmembrane membrane proteins found in almost all mammalian cell types. P2 receptors can be further divided into two subclasses, P2X (ionotropic receptors, P2XRs, members of ligand-gated ion channels) and P2Y (metabotropic receptors, P2YRs, members of G protein-coupled receptors) ^14^ Using Ca^2+^-imaging with a red fluorescence Ca^2+^ indicator (XRhod1-AM) in NIH-3T3 cells, we found that a 1-min 100-μM ATP application elicited a surge of cytosolic Ca^2+^ and the same ATP stimulus 1 min later generated a second Ca^2+^-response smaller than the first. Both responses were blocked by Suramin (antagonist for P2X and P2Y receptors) (Fig. S6a), suggesting that P2 receptors are the major purinergic receptors mediating this ATP-induced intracellular Ca^2+^ increase.

Although both can be activated by extracellular ATP, P2XRs and P2YRs produce cytosolic Ca^2+^ surges via different pathways: P2XRs are ionotropic receptors, allowing the influx of extracellular Ca^2+^ upon ATP-binding, and P2YRs are metabotropic receptors that trigger the release of Ca^2+^ from internal stores after ATP-binding ^15^. The fact that the second Ca^2+^-response was always smaller than the first suggested that the source of Ca^2+^ was unlikely extracellular Ca^2+^ (external [Ca^2+^] is constant and the reactivation time for P2XRs is much shorter than 1 minute) but rather internal Ca^2+^ stores, which require more than a minute to be fully refilled ^16^. This notion is supported by the following empirical evidence. When we prohibited the refill of internal Ca^2+^ stores by removing extracellular Ca^2+^ using 0 Ca^2+^/EGTA Tyrode’s solution, the second ATP-induced Ca^2+^-response diminished (Fig. S6a), suggesting a P2YR-mediated pathway involving internal Ca^2+^-store. Furthermore, the fact that ATP failed to evoke any detectable inward current (Fig. S6b) disputes the involvement of ionotropic P2XRs in 3T3 cells. Thus, we conclude that P2YRs are the dominant mediator for the ATP-induced Ca^2+^ response in 3T3 cells.

### Supplementary Figures and Legends

**Figure S1.**
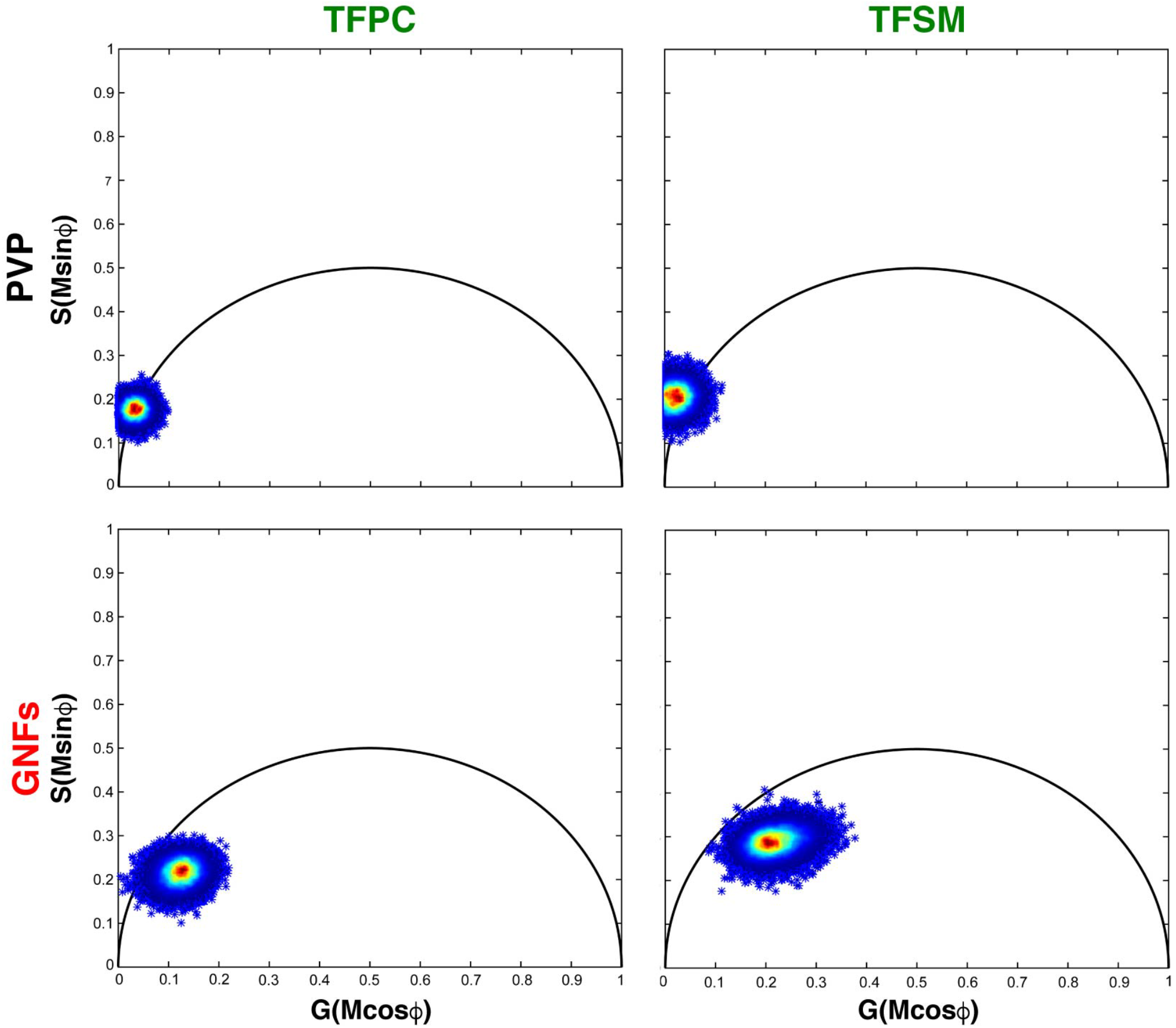
Fluorescence lifetimes of TFPC and TFSM are minimally affected by GNFs. Phasor plots of TFPC and TFSM mixed with PVP or GNFs. TFPC, TopFluor Phosphocholine, TFSM, TopFluor Sphingomyelin.

**Figure S2.**
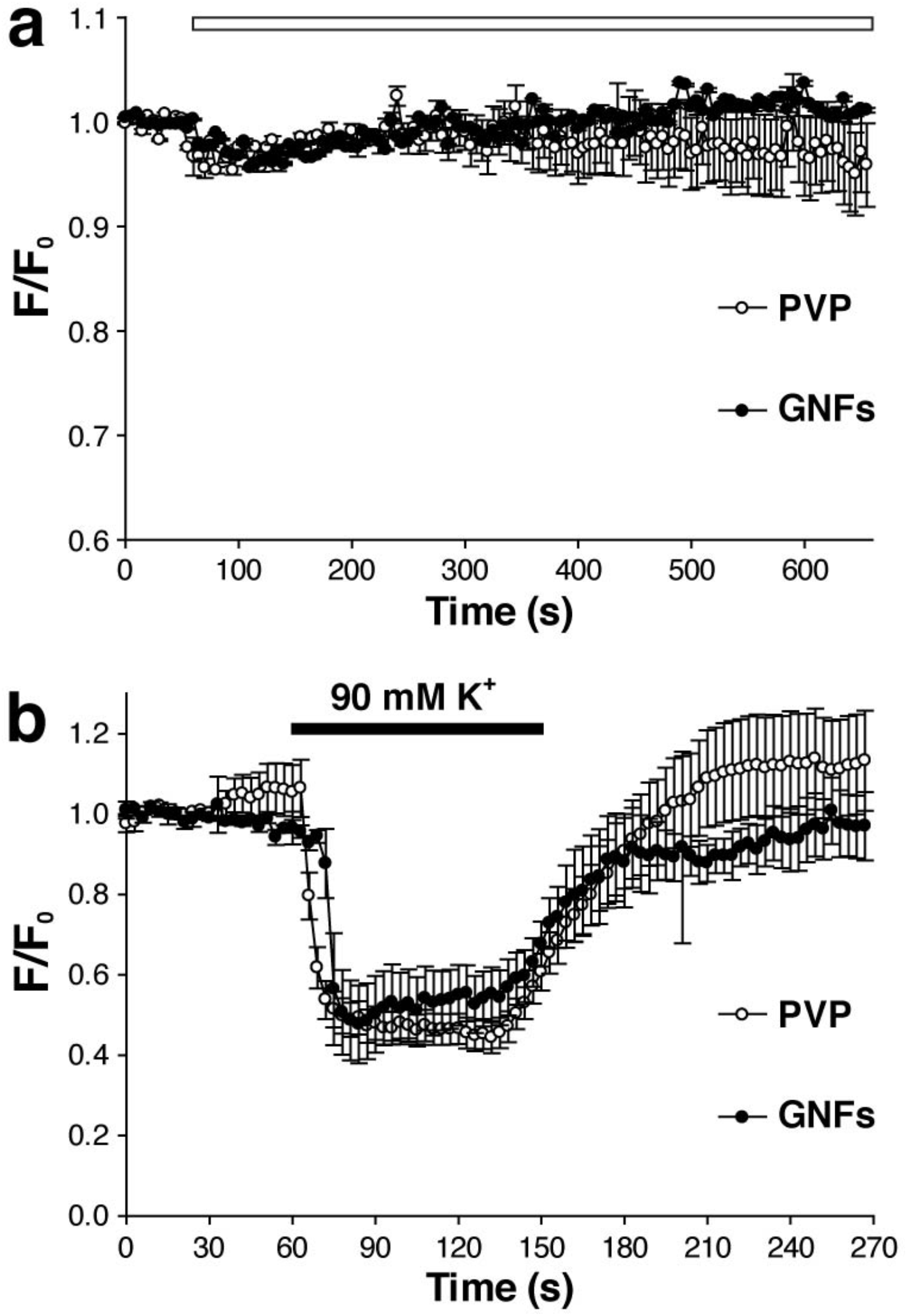
Acute application of GNFs does not affect DiO fluorescence or its voltage sensing capabilities. **(a)** Effect of PVP or GNFs (white bar, starting at 60s and lasting 600s) on DiO fluorescence (both n = 5 replicates; *p* > 0.1). **(b)** Decrease in DiO fluorescence (when paired with DPA) induced by 90 mM K^+^ for 5 min application of PVP or GNFs (both n = 5 replicates; *p* > 0.05).

**Figure S3.**
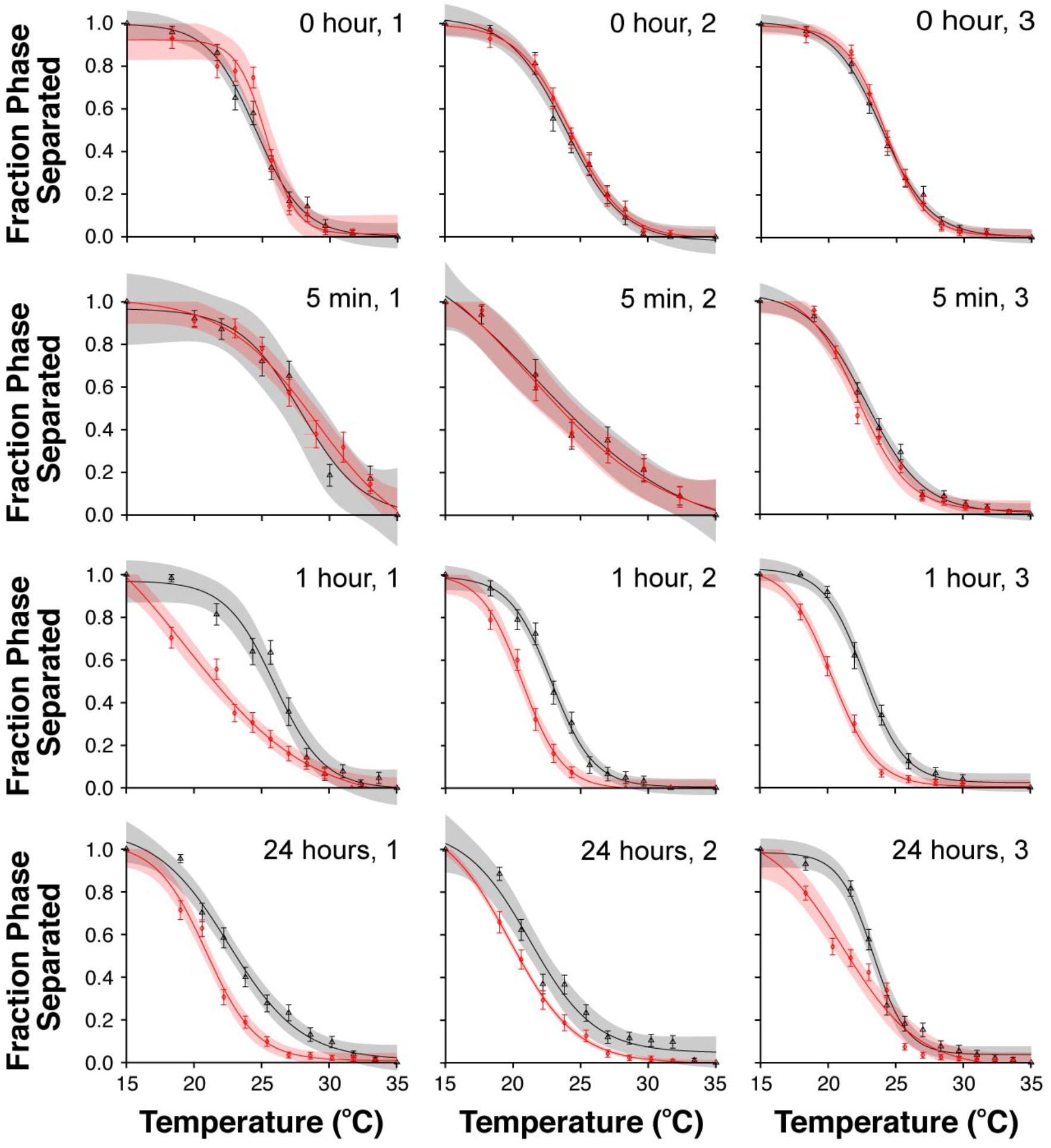
Application of GNFs prior to GPMV isolation reduces miscibility temperature. Phase separation of 3T3 cells pre-incubated with media containing either PVP (black) or GNFs (red). Phase separated fractions were calculated from the total numbers of phase-separated and non-separated GPMVs at each temperature point. Areas of light shading show the 95% confidence interval on the sigmoidal fit function, defined as: 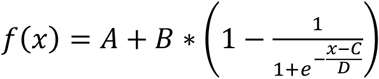.

**Figure S4.**
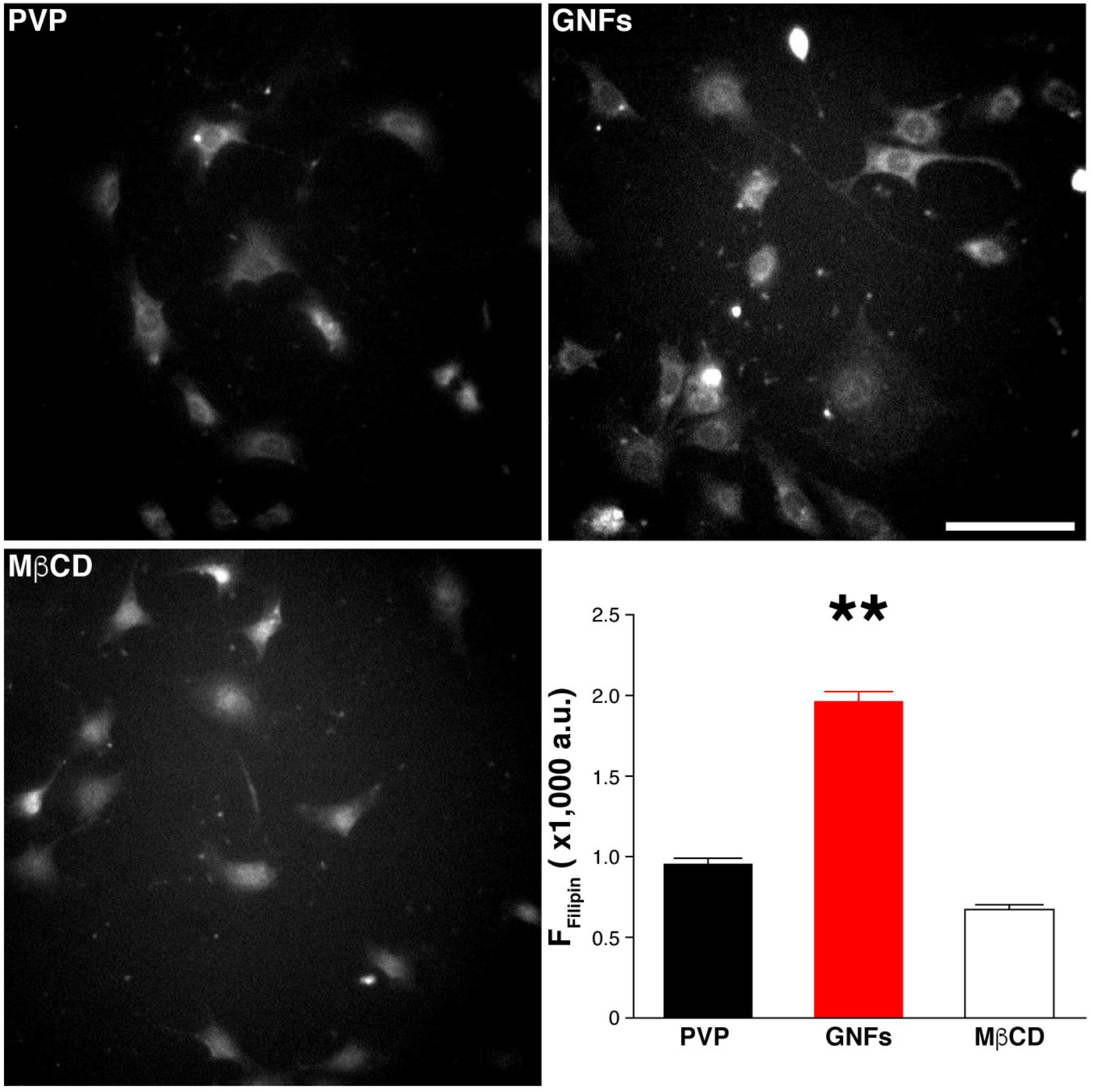
Filipin staining shows GNF-induced cell surface cholesterol increase. Scale bar, 100 μm; n = 9 replicates. \*\**p* < 0.01. Error bars represent the S.E.M.

**Figure S5.**
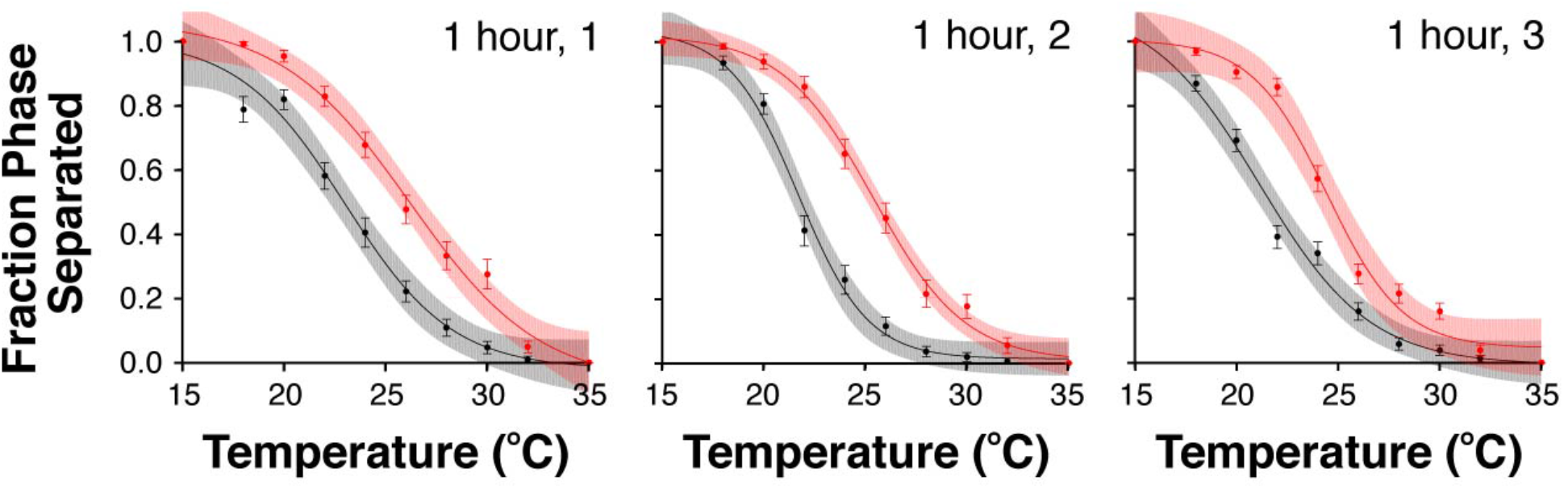
1-hour application of GNFs after GPMV isolation increases miscibility temperature. Phase separation of isolated GPMVs treated with either PVP (black) or GNFs (red) for 1 h. Phase separated fractions were calculated from the total numbers of phase-separated and non-separated GPMVs at each temperature point. Areas of light shading show the 95% confidence interval on the sigmoidal fit function, defined as: 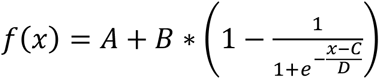.

**Figure S6.**
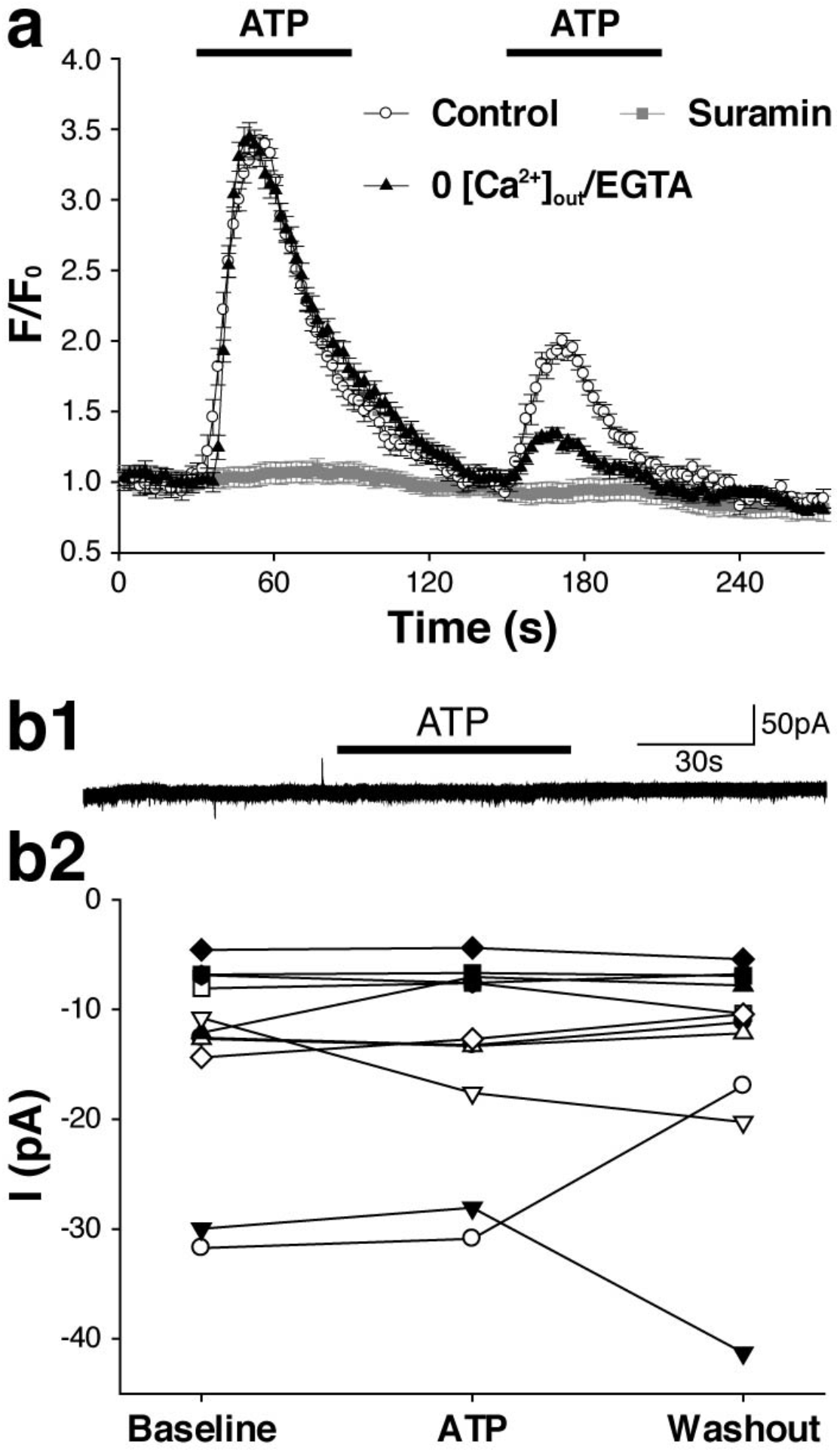
P2YRs mediate ATP-induced Ca^2+^-release from internal Ca^2+^ -store. **(a)** Average Ca^2+^-response upon 100μM ATP applications in Tyrode’s solution or Tyrode’s containing Suramin (100 μM) or 0 Ca^2+^/EGTA (2 mM). **(b1)** Sample record of a 3T3 cell before, during, and after 100 μM ATP challenge. **(b2)** Average currents within the 10 s period of 3T3 cells before, during, and after 100 μM ATP challenge (n = 11 replicates; *p* > 0.05).

**Figure S7.**
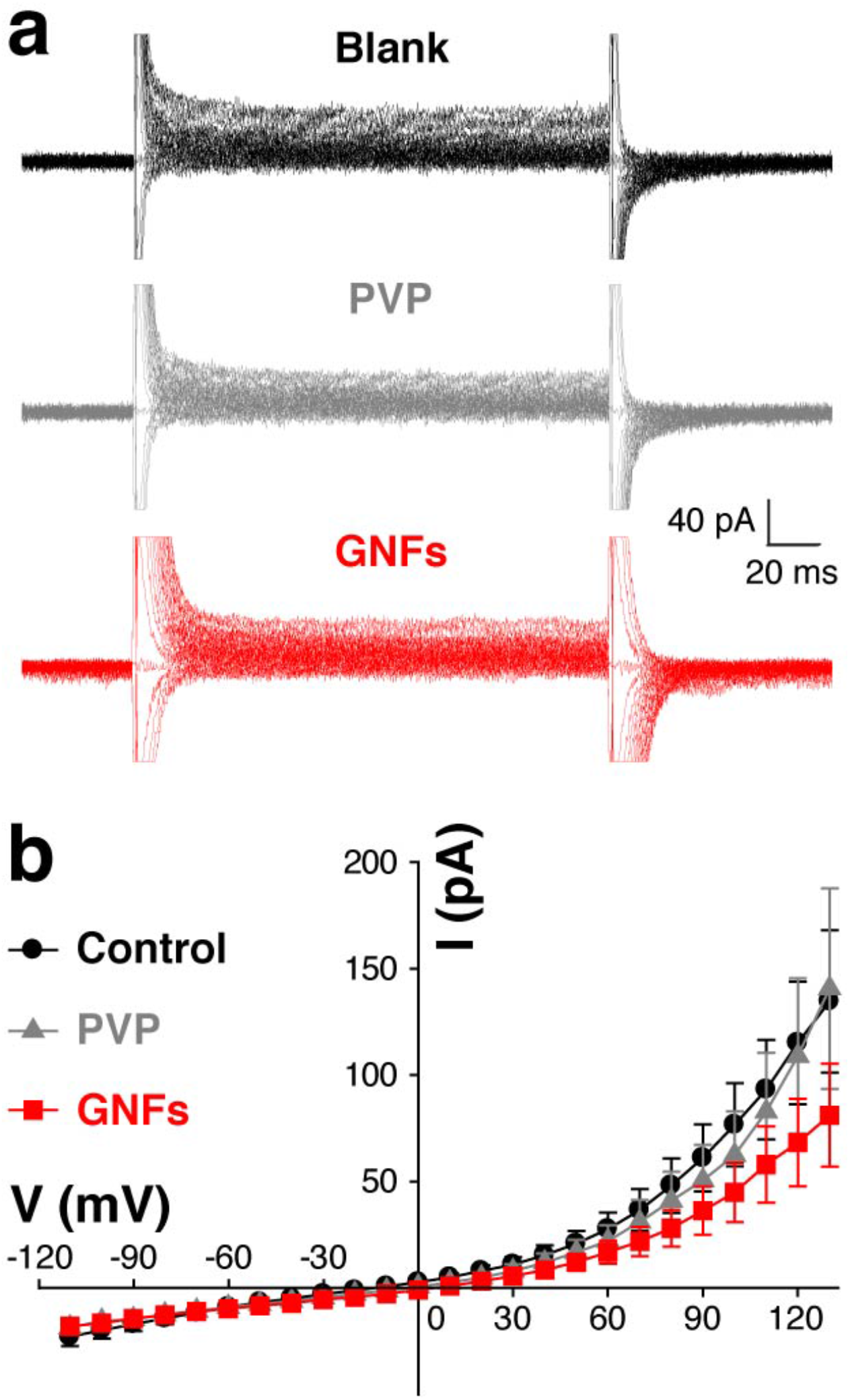
Acute application of GNFs does not alter current-voltage characteristics in 3T3 cells. **(a)** Trace overlays for a 3T3 cell in Tyrode or Tyrode containing either PVP or GNFs. **(b)** Average I-V curves of the same 3T3 cells subjected to three treatments (n = 5 replicates; *p* > 0.05).

**Figure S8.**
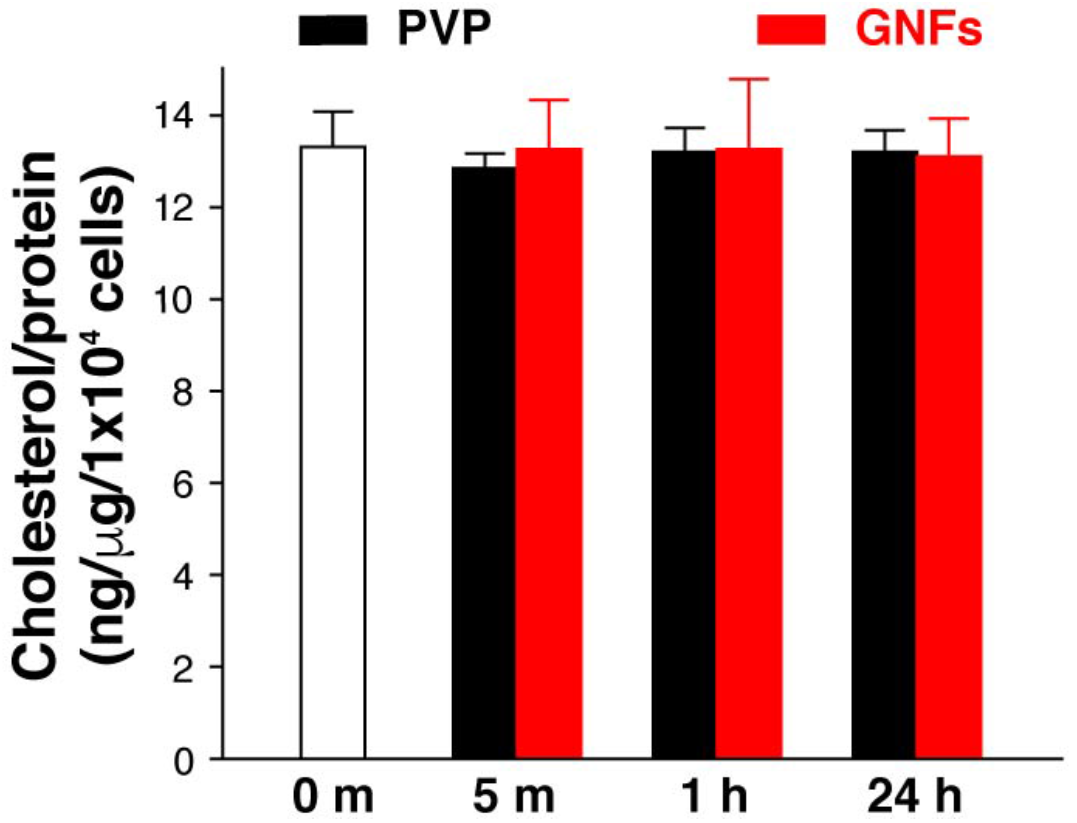
GNFs do not change total cellular cholesterol. Total cellular cholesterol quantifications by gas chromatography coupled with a flame ionization detector after specified periods of incubation with PVP or GNFs (all n = 6 replicates; all *p* > 0.05).

**Figure S9.**
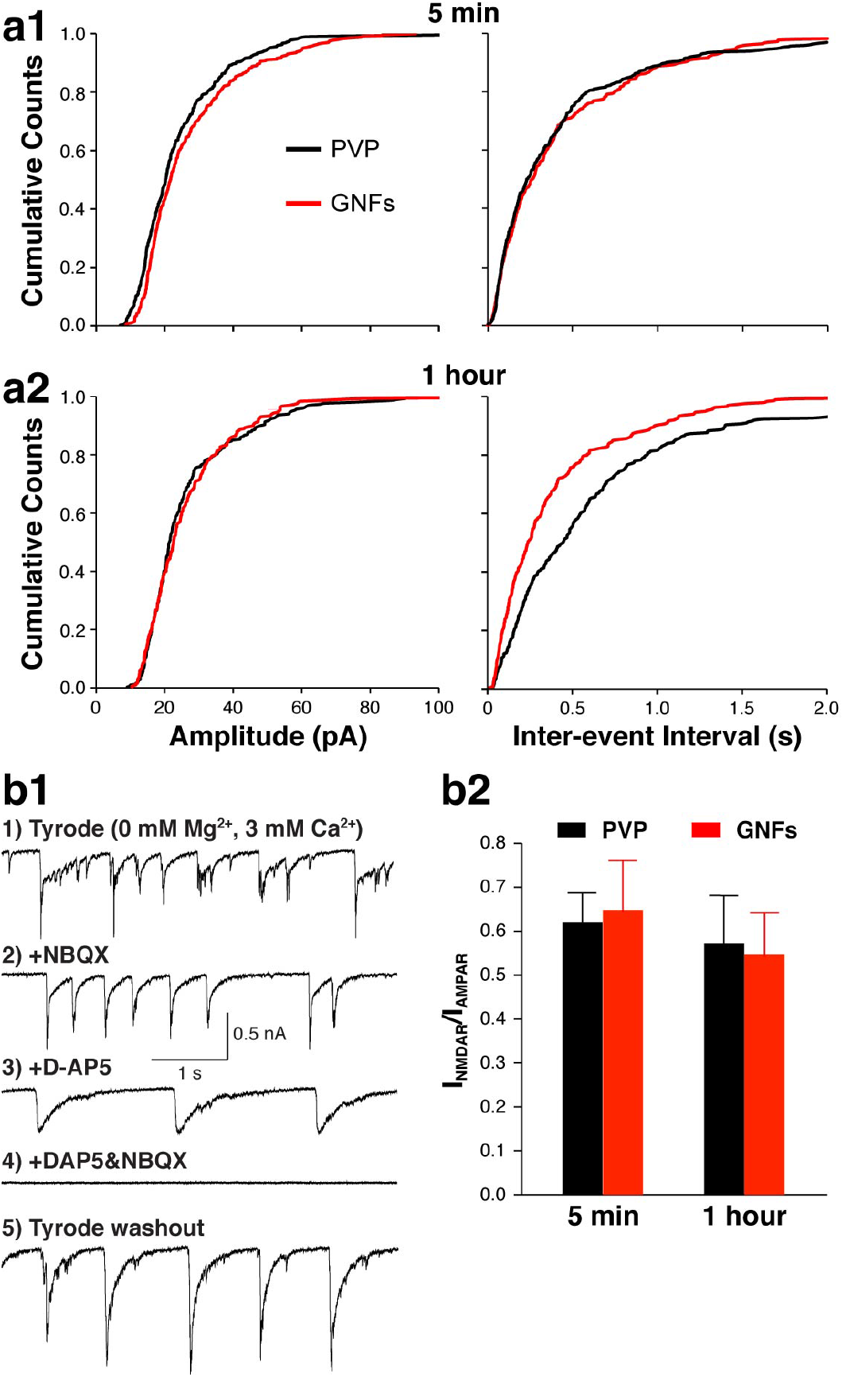
Acute treatment with GNFs has minimal postsynaptic effect on neurons. **(a)** Cumulative distributions of mEPSC amplitudes and frequencies in neurons treated with PVP or GNFs for 5 min or 1 h (5 min, n_PVP_ = 8 replicates, n_graphene_ = 8 replicates, 1 h, n_PVP_ = 6 replicates, n_graphene_ = 6 replicates; *p* < 0.01 between 1-hour PVP and GNF treatment, *p* > 0.05 for all other conditions). (b1) Sequential recording of AMPA and NMDA receptor currents from the same neurons. (b2) NMDAR and AMPAR current ratio in PVP or GNFs treated neurons (5 min, nPVP = 11 replicates, ngraphene = 12 replicates, 1 h, nPVP = 10 replicates, ngraphene = 9 replicates; all *p* > 0.05).

**Figure S10.**
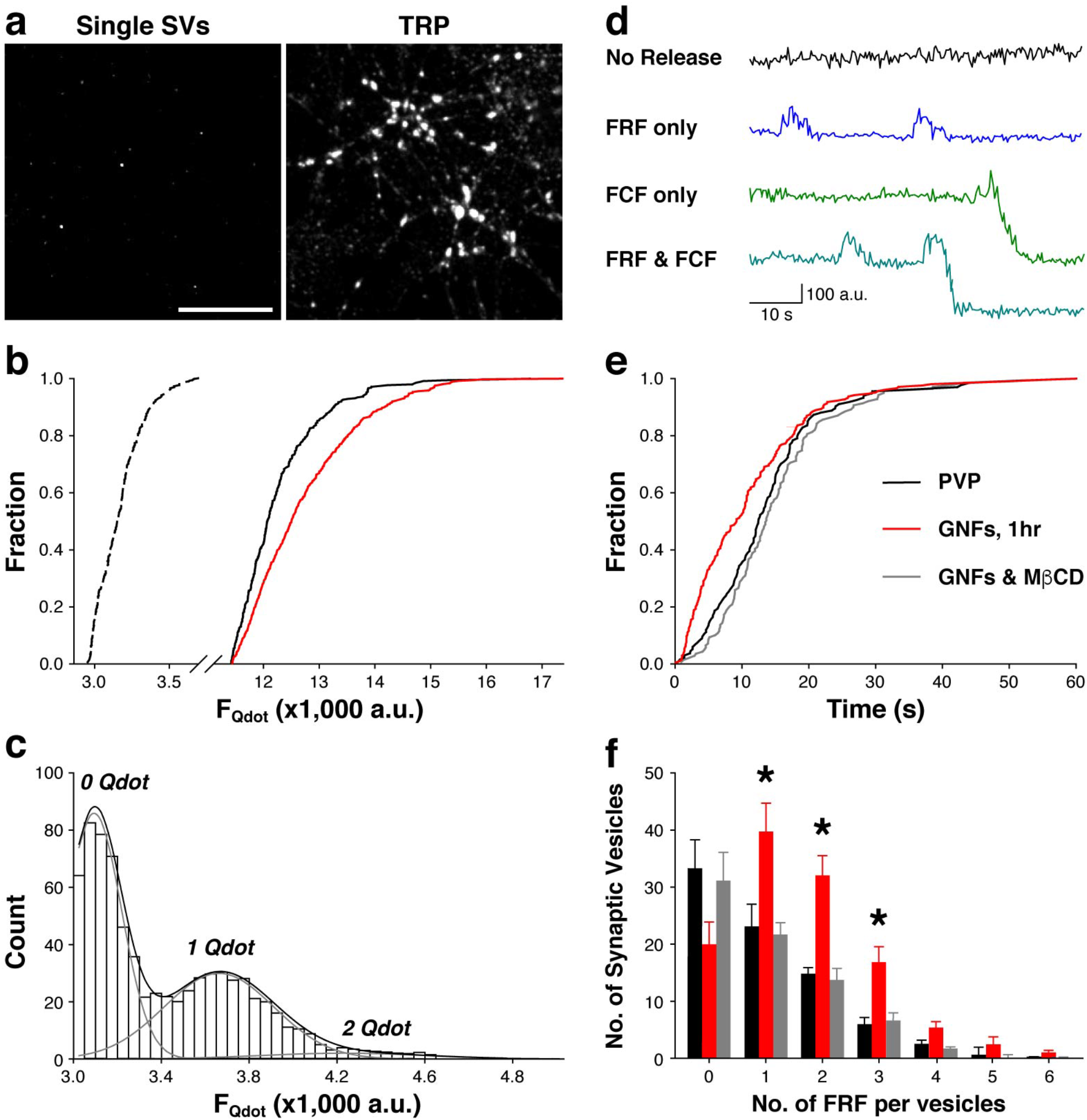
Single Qdot imaging reports single vesicle fusion kinetics. TRP, total releasable pool, FRF, fast and reversible fusion, FCF, full collapse fusion. **(a)** Sample images of single synaptic vesicle loading for releasable pool (0.8 nM Qdot) or TRP loading (100 nM Qdot). **(b)** Synaptic Qdot photoluminescence intensities from single vesicle and TRP loading were measured with the same settings and plotted on the same scale. The background intensity was 2,784 ± 96 a.u. and the detection threshold was set at 2,900 a.u.. Under single vesicle loading, the mean Qdot photoluminescence intensity was 402 ± 43 a.u.. Under total recycling pool loading, the mean Qdot photoluminescence intensities were 8,913 ± 144 a.u. for neurons treated with PVP and 9,980 ± 185 a.u. for neurons treated with graphene, which is significantly higher *(p* < 0.05). Based on these intensity values, we estimated that the average numbers of total recycling vesicles are 23.4 for neurons on glass and 26.2 for those on graphene. **(c)** The distribution of mean Qdot photoluminescence in individual synaptic boutons defined by retrospective FM4-64 staining. Quantal analysis (black and gray lines) indicates that the mean photoluminescence intensity of loaded single Qdots was 388 ± 71 a.u. after background subtraction. **(d)** Sample photoluminescence traces of single Qdots for four different types of synaptic vesicle release behaviors. (e) Synaptic vesicle release probability (measured as the first fusion event of Qdot-loaded synaptic vesicles) in neurons treated with PVP, GNFs or GNFs & MβCD (GNFs 1 hr, *p* < 0.05). (f) Distribution of individual synaptic vesicles conducting different rounds of FRF. \**p* < 0.05. Error bars represent the S.E.M.

