## Supplemental Materials for "Graphene nanoflakes for acute manipulation of membrane cholesterol and transmembrane signaling"

**Supplementary Figures and Legends:**

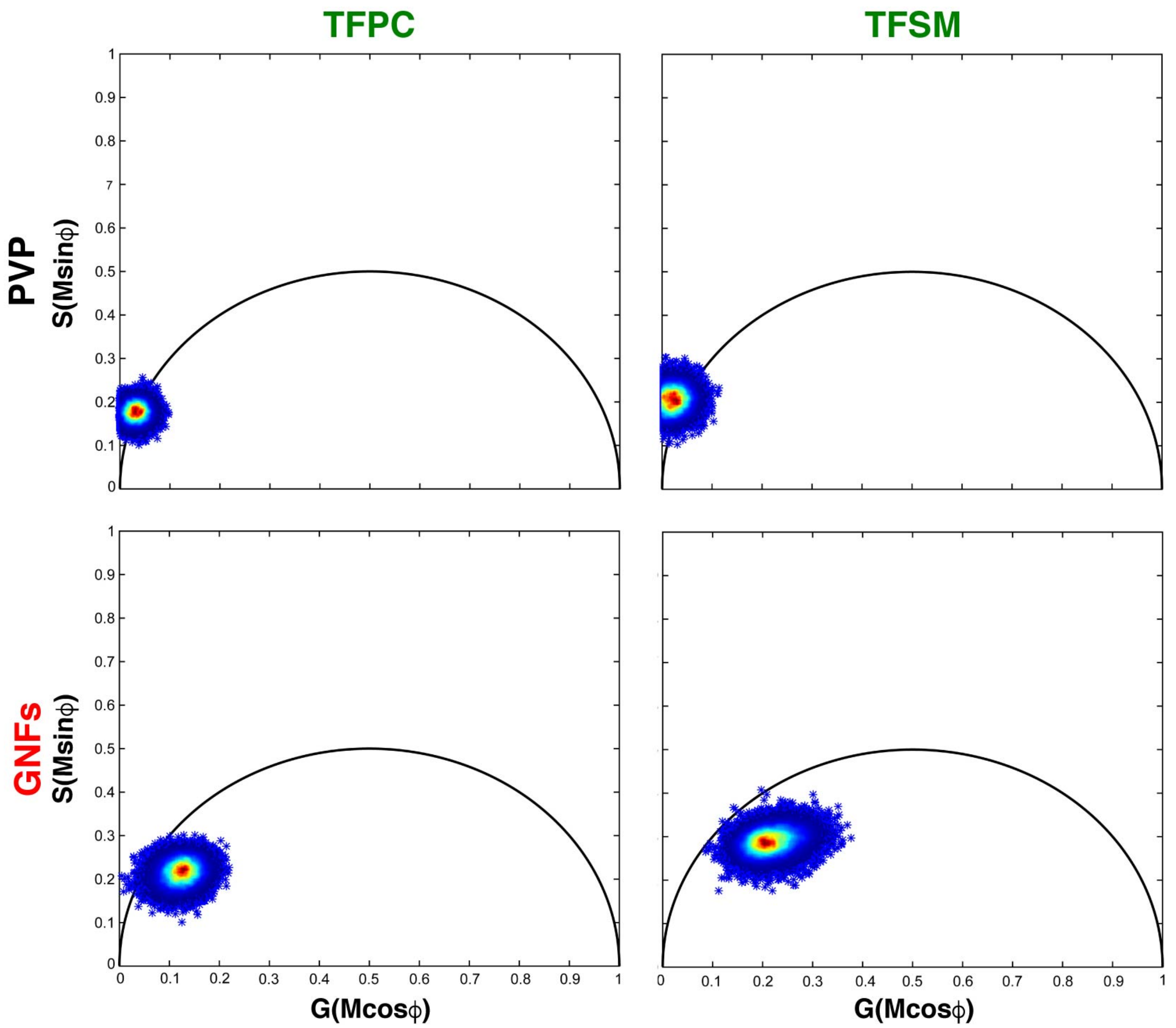


**Figure S1. Fluorescence lifetimes of TFPC and TFSM are minimally affected by GNFs.** Phasor plots of TFPC and TFSM mixed with PVP or GNFs. TFPC, TopFluor Phosphocholine, TFSM, TopFluor Sphingomyelin.

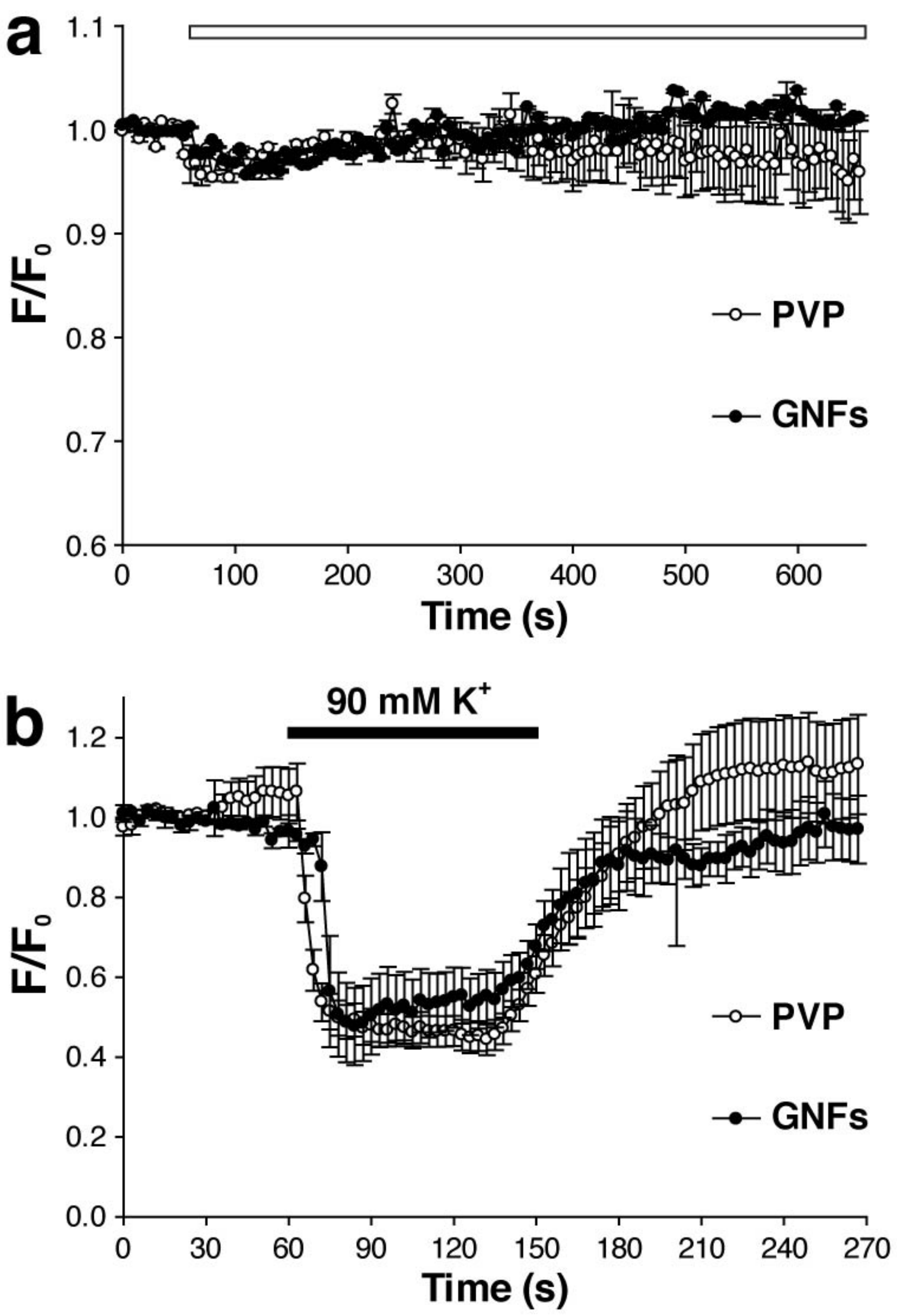


**Figure S2. Acute application of GNFs does not affect DiO fluorescence or its voltage sensing capabilities. (a)** Effect of PVP or GNFs (white bar, starting at 60s and lasting 600s) on DiO fluorescence (both n = 5 replicates; $p > 0.1$). **(b)** Decrease in DiO fluorescence (when paired with DPA) induced by 90 mM $K^+$ for 5 min application of PVP or GNFs (both n = 5 replicates; $p > 0.05$).

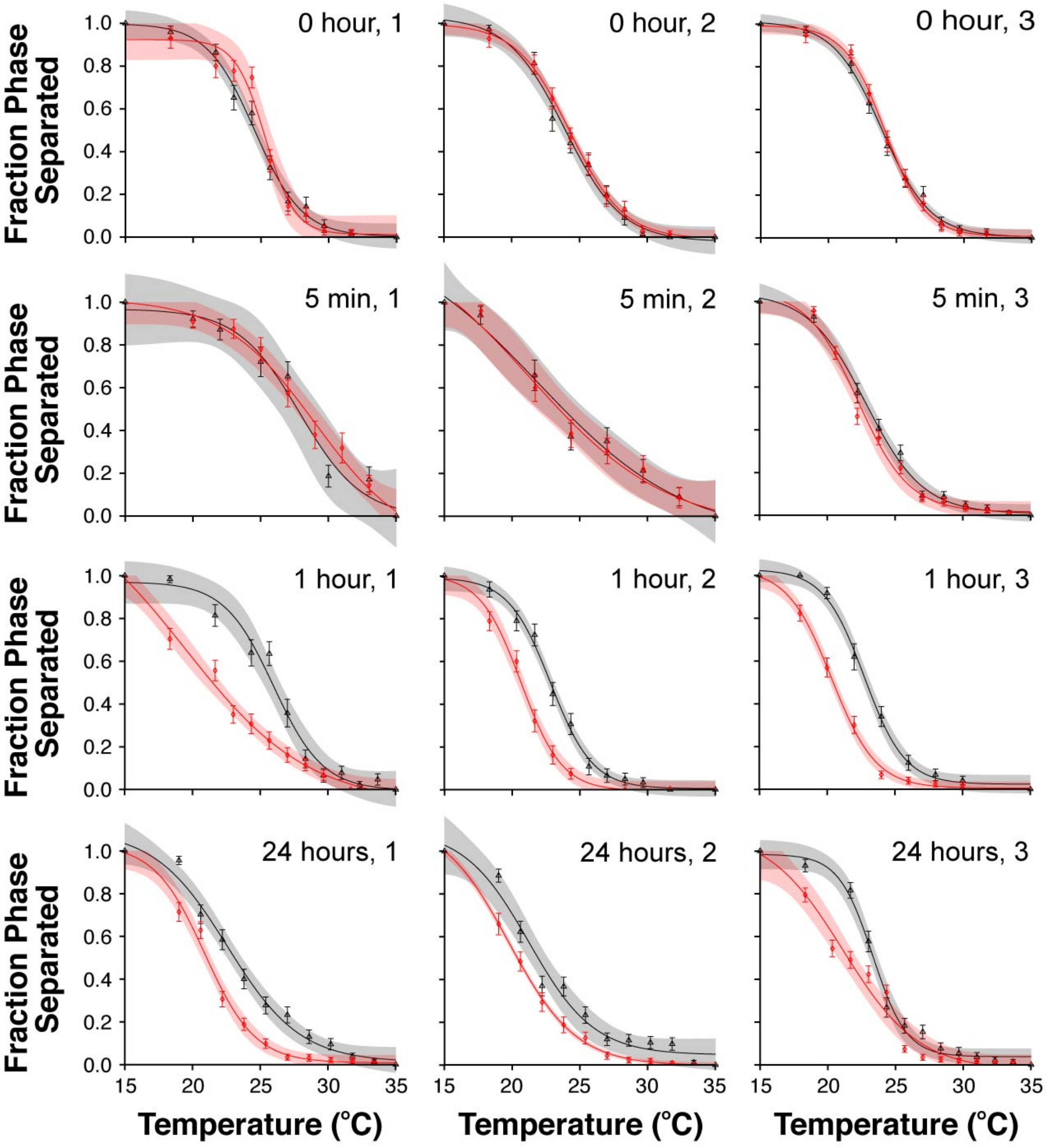


**Figure S3. Application of GNFs prior to GPMV isolation reduces miscibility temperature.** Phase separation of 3T3 cells pre-incubated with media containing either PVP (black) or GNFs (red). Phase separated fractions were calculated from the total numbers of phase-separated and non-separated GPMVs at each temperature point. Areas of light shading show the 95% confidence interval on the sigmoidal fit function, defined as: $f(x) = A + B * \left(1 - \frac{1}{1+e^{-\frac{x-C}{D}}}\right)$.

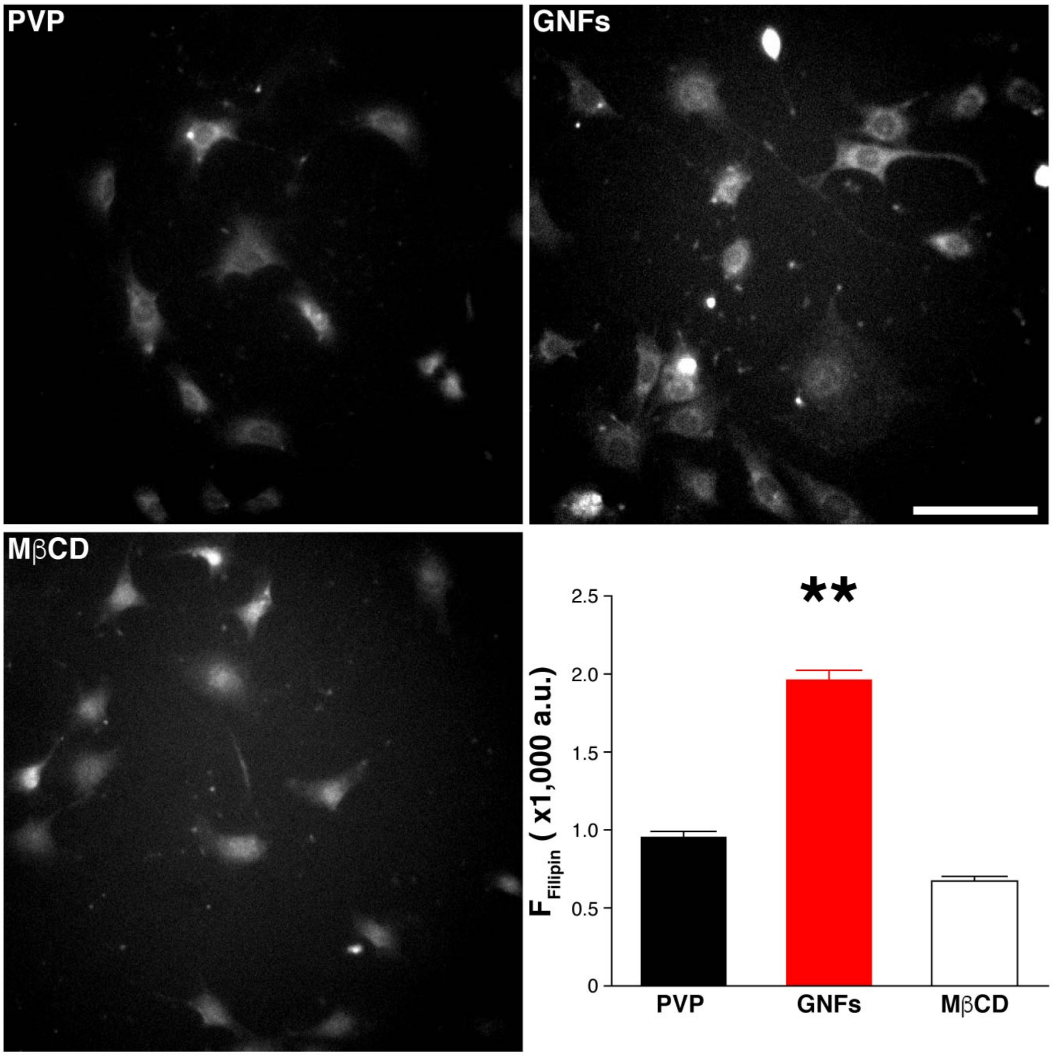


**Figure S4. Filipin staining shows GNF-induced cell surface cholesterol increase.** Scale bar, 100 μm; n = 9 replicates. ***p* < 0.01. Error bars represent the S.E.M.

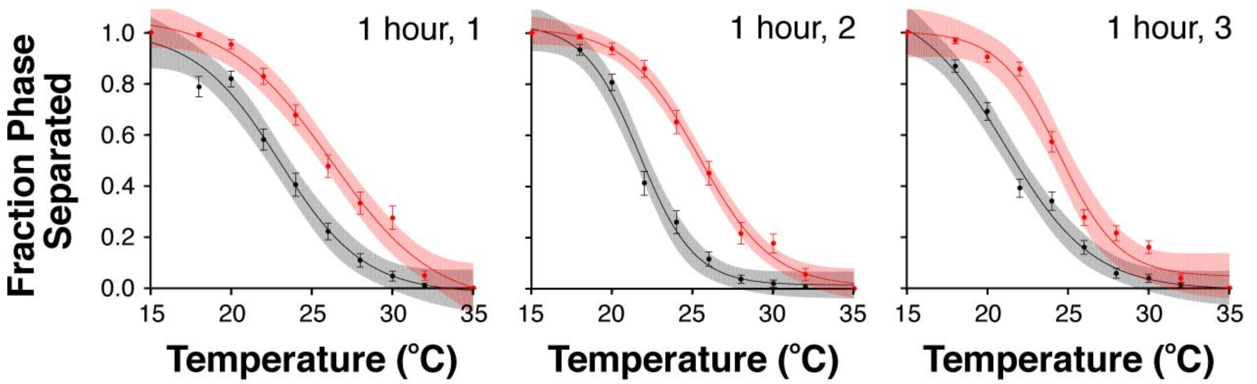


**Figure S5. 1-hour application of GNFs after GPMV isolation increases miscibility temperature.** Phase separation of isolated GPMVs treated with either PVP (black) or GNFs (red) for 1 h. Phase separated fractions were calculated from the total numbers of phase-separated and non-separated GPMVs at each temperature point. Areas of light shading show the 95% confidence interval on the sigmoidal fit function, defined as: $f(x) = A + B * \left(1 - \frac{1}{1+e^{-\frac{x-C}{D}}}\right)$.

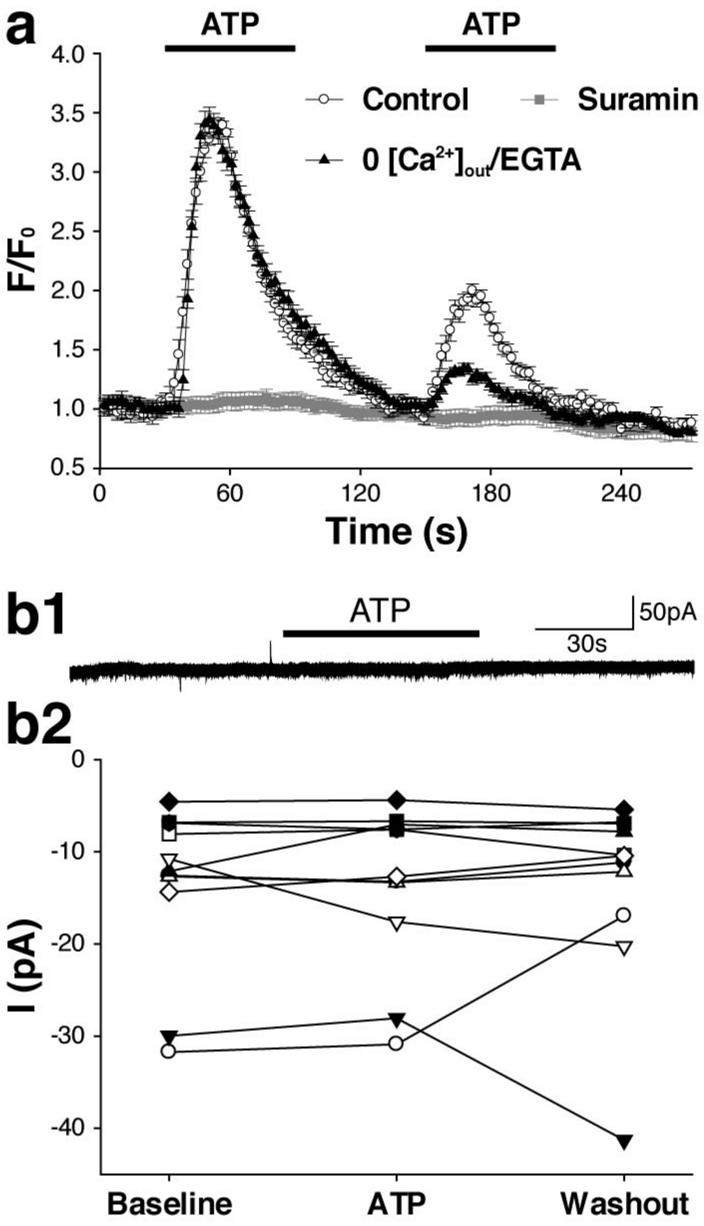


**Figure S6. P2YRs mediate ATP-induced $Ca^{2+}$-release from internal $Ca^{2+}$-store. (a)** Average $Ca^{2+}$-response upon 100μM ATP applications in Tyrode's solution or Tyrode's containing Suramin (100 μM) or 0 $Ca^{2+}$/EGTA (2 mM). **(b1)** Sample record of a 3T3 cell before, during, and after 100 μM ATP challenge. **(b2)** Average currents within the 10 s period of 3T3 cells before, during, and after 100 μM ATP challenge (n = 11 replicates; $p > 0.05$).

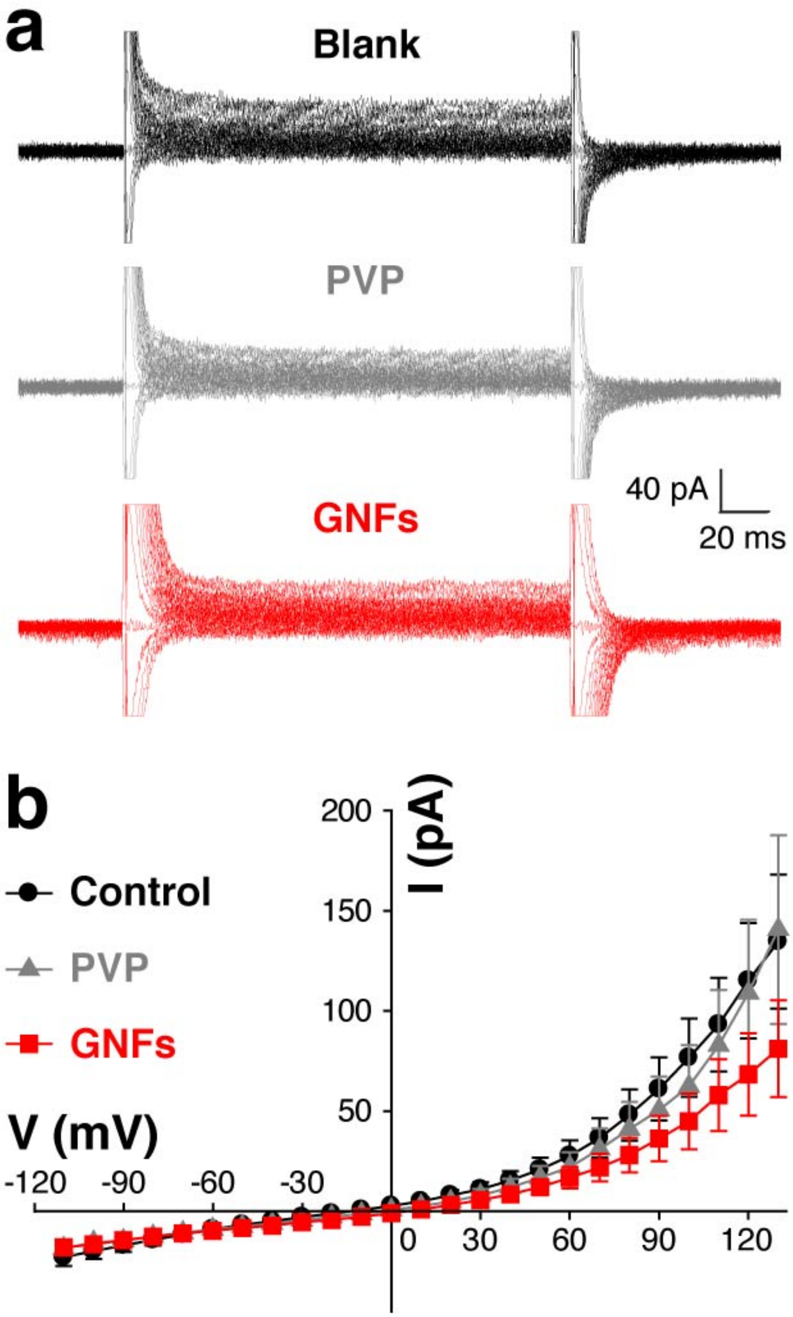


**Figure S7. Acute application of GNFs does not alter current-voltage characteristics in 3T3 cells. (a)** Trace overlays for a 3T3 cell in Tyrode or Tyrode containing either PVP or GNFs. **(b)** Average I-V curves of the same 3T3 cells subjected to three treatments (n = 5 replicates; $p > 0.05$).

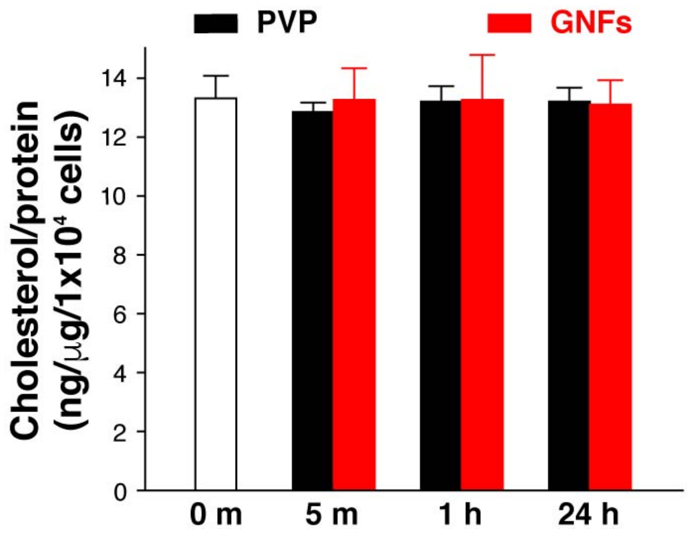


**Figure S8. GNFs do not change total cellular cholesterol.** Total cellular cholesterol quantifications by gas chromatography coupled with a flame ionization detector after specified periods of incubation with PVP or GNFs (all n = 6 replicates; all $p > 0.05$).

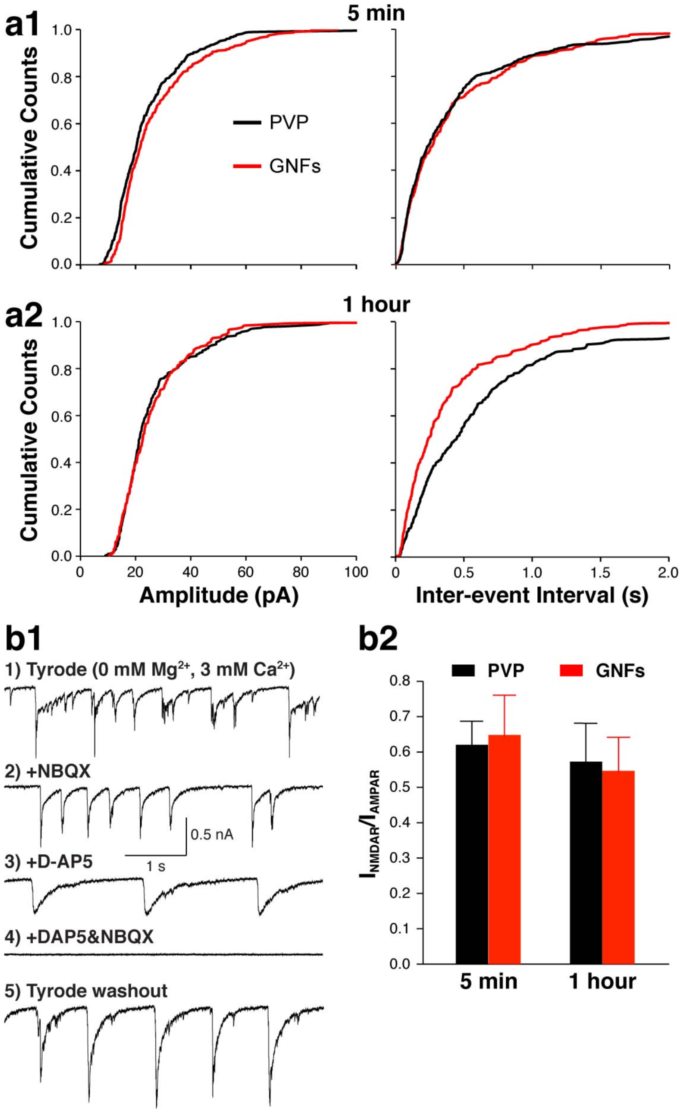


**Figure S9. Acute treatment with GNFs has minimal postsynaptic effect on neurons.** (**a**) Cumulative distributions of mEPSC amplitudes and frequencies in neurons treated with PVP or GNFs for 5 min or 1 h (5 min, $n_{PVP}$ = 8 replicates, $n_{graphene}$ = 8 replicates, 1 h, $n_{PVP}$ = 6 replicates, $n_{graphene}$ = 6 replicates; $p < 0.01$ between 1-hour PVP and GNF treatment, $p > 0.05$ for all other conditions). (**b1**) Sequential recording of AMPA and NMDA receptor currents from the same neurons. (**b2**) NMDAR and AMPAR current ratio in PVP or GNFs treated neurons (5 min, $n_{PVP}$ = 11 replicates, $n_{graphene}$ = 12 replicates, 1 h, $n_{PVP}$ = 10 replicates, $n_{graphene}$ = 9 replicates; all $p > 0.05$).

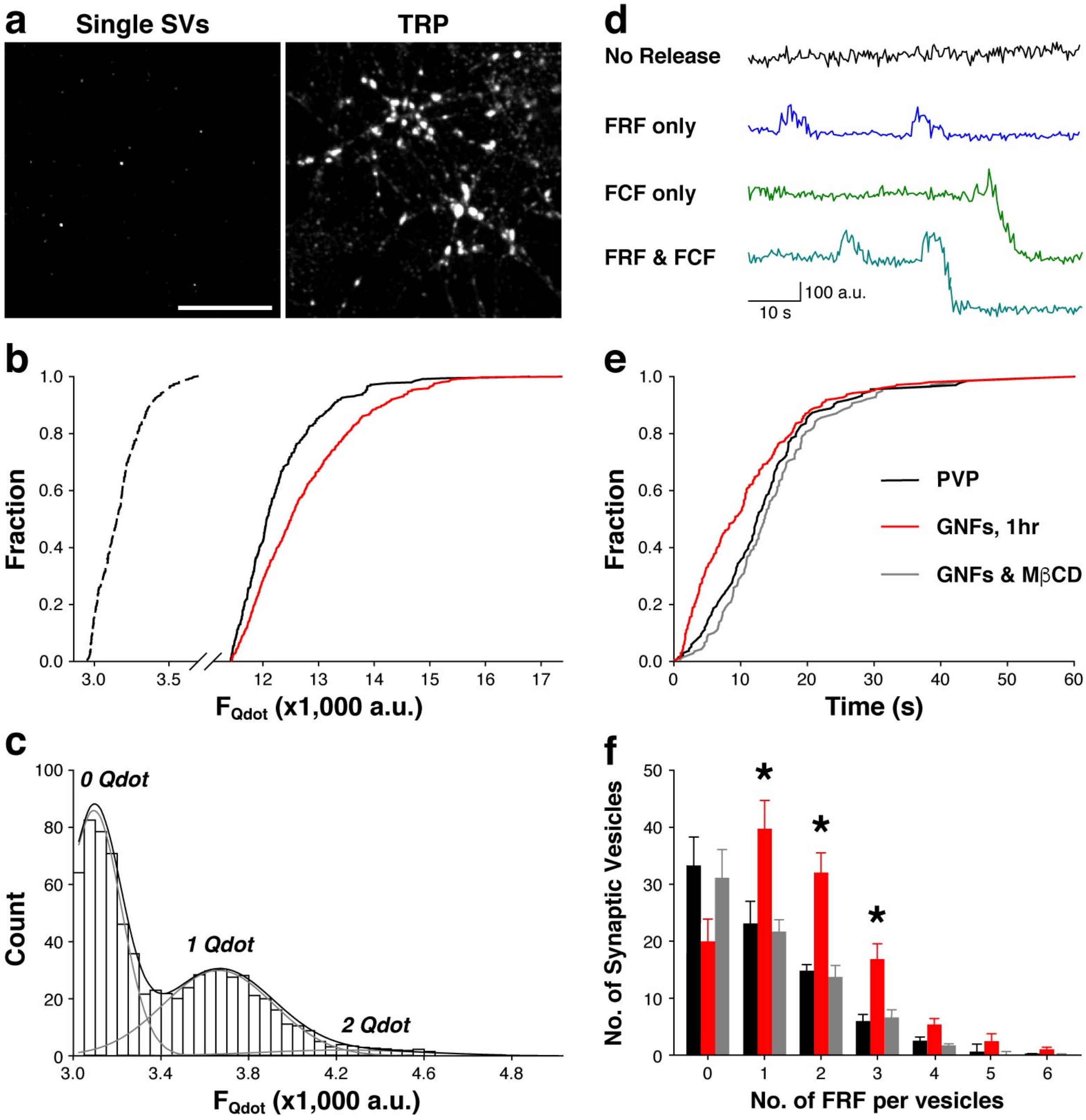


**Figure S10. Single Qdot imaging reports single vesicle fusion kinetics.** TRP, total releasable pool, FRF, fast and reversible fusion, FCF, full collapse fusion. (**a**) Sample images of single synaptic vesicle loading for releasable pool (0.8 nM Qdot) or TRP loading (100 nM Qdot). (**b**) Synaptic Qdot photoluminescence intensities from single vesicle and TRP loading were measured with the same settings and plotted on the same scale. The background intensity was 2,784 ± 96 a.u. and the detection threshold was set at 2,900 a.u.. Under single vesicle loading, the mean Qdot photoluminescence intensity was 402 ± 43 a.u.. Under total recycling pool loading, the mean Qdot photoluminescence intensities were 8,913 ± 144 a.u. for neurons treated with PVP and 9,980 ± 185 a.u. for neurons treated with graphene, which is significantly higher ($p < 0.05$). Based on these intensity values, we estimated that the average numbers of total recycling vesicles are 23.4 for neurons on glass and 26.2 for those on graphene. (**c**) The distribution of mean Qdot photoluminescence in individual synaptic boutons defined by retrospective FM4-64 staining. Quantal analysis (black and gray lines) indicates that the mean photoluminescence intensity of loaded single Qdots was 388 ± 71 a.u. after background subtraction. (**d**) Sample photoluminescence traces of single Qdots for four different types of synaptic vesicle release behaviors. (**e**) Synaptic vesicle release probability (measured as the first fusion event of Qdot-loaded synaptic vesicles) in neurons treated with PVP, GNFs or GNFs & MβCD (GNFs 1 hr, $p < 0.05$). (**f**) Distribution of individual synaptic vesicles conducting different rounds of FRF. *$p < 0.05$. Error bars represent the S.E.M.
